# SensitiveCancerGPT: Leveraging Generative Large Language Model on Structured Omics Data to Optimize Drug Sensitivity Prediction

**DOI:** 10.1101/2025.02.27.640661

**Authors:** Shaika Chowdhury, Sivaraman Rajaganapathy, Lichao Sun, Liewei Wang, Ping Yang, James R Cerhan, Nansu Zong

## Abstract

**Objective:** The fast accumulation of vast pharmacogenomics data of cancer cell lines provide unprecedented opportunities for drug sensitivity prediction (DSP), a crucial prerequisite for the advancement of precision oncology. Recently, Generative Large Language Models (LLM) have demonstrated performance and generalization prowess across diverse tasks in the field of natural language processing (NLP). However, the structured format of the pharmacogenomics data poses challenge for the utility of LLM in DSP. Therefore, the objective of this study is multi-fold: to adapt prompt engineering for structured pharmacogenomics data toward optimizing LLM’s DSP performance, to evaluate LLM’s generalization in real-world DSP scenarios, and to compare LLM’s DSP performance against that of state-of-the-science baselines.

**Methods:** We systematically investigated the capability of the Generative Pre-trained Transformer (GPT) as a DSP model on four publicly available benchmark pharmacogenomics datasets, which are stratified by five cancer tissue types of cell lines and encompass both oncology and non-oncology drugs. Essentially, the predictive landscape of GPT is assessed for effectiveness on the DSP task via four learning paradigms: zero-shot learning, few-shot learning, fine-tuning and clustering pretrained embeddings. To facilitate GPT in seamlessly processing the structured pharmacogenomics data, domain-specific novel prompt engineering is employed by implementing three prompt templates (i.e., Instruction, Instruction-Prefix, Cloze) and integrating pharmacogenomics-related features into the prompt. We validated GPT’s performance in diverse real-world DSP scenarios: cross-tissue generalization, blind tests, and analyses of drug-pathway associations and top sensitive/resistant cell lines. Furthermore, we conducted a comparative evaluation of GPT against multiple Transformer-based pretrained models and existing DSP baselines.

**Results:** Extensive experiments on the pharmacogenomics datasets across the five tissue cohorts demonstrate that fine-tuning GPT yields the best DSP performance (28% F1 increase, p-value= 0.0003) followed by clustering pretrained GPT embeddings (26% F1 increase, p-value= 0.0005), outperforming GPT in-context learning (i.e., few-shot). However, GPT in the zero-shot setting had a big F1 gap, resulting in the worst performance. Within the scope of prompt engineering, performance enhancement was achieved by directly instructing GPT about the DSP task and resorting to a concise context format (i.e., instruction-prefix), leading to F1 performance gain of 22% (p-value=0.02); while incorporation of drug-cell line prompt context derived from genomics and/or molecular features further boosted F1 score by 2%. Compared to state-of-the-science DSP baselines, GPT significantly asserted superior mean F1 performance (16% gain, p-value<0.05) on the GDSC dataset. In the crosstissue analysis, GPT showcased comparable generalizability to the within-tissue performances on the GDSC and PRISM datasets, while statistically significant F1 performance improvements on the CCLE (8%, p-value=0.001) and DrugComb (19%, p-value=0.009) datasets. Evaluation on the challenging blind tests suggests GPT’s competitiveness on the CCLE and DrugComb datasets compared to random splitting. Furthermore, analyses of the drug-pathway associations and log probabilities provided valuable insights that align with previous DSP findings.

**Conclusion:** The diverse experiment setups and in-depth analysis underscore the importance of generative LLM, such as GPT, as a viable in silico approach to guide precision oncology.

**Availability:** https://github.com/bioIKEA/SensitiveCancerGPT

## 1. Introduction

Cancer is a complex genetic disease that originates from the accumulation of gene mutations within a cell and is ranked as the second leading cause of death in the United States, according to the American Cancer Society (Siegel et al. 2023). Given the tumor heterogeneity arising from the genetic variations among patients even with the same cancer type (Bedard et al. 2013), substantial differences in the anti-cancer drug response can be expected, thereby highlighting the urgent need for targeted therapies. Owing to the high cost and time associated with developing and validating anti-cancer drugs in clinical trials - which is further exacerbated by the 96% failure rate (Kola et al. 2004) – it is imperative to develop preclinical computational models that can accurately predict whether a cell line is sensitive or resistant to a particular drug. The availability of genomic profiles of cancer cell lines (Barretina et al. 2012, Yang et al. 2012) collected via high-throughput screening technologies offers feasible resources to develop robust drug response models and identify the important biomarkers predictive of drug sensitivity.

The advent of cutting-edge machine learning methods, especially deep learning-based ones, coupled with large-scale pharmacogenomics datasets, has ushered in the development of data-driven computational methods for the effective prediction of cancer drug response (Li et al. 2019, Chiu et al. 2019, Zuo et al. 2021, Manica et al. 2019 and G et al. 2020) (Literature Review presented in **Supplementary Note 1**). These methods formalize DSP either into a regression or classification problem contingent on the prediction of a continuous (i.e., IC50) or categorical (i.e., sensitive or resistant) output, respectively. Modeling this problem involves feature extraction from the drug’s molecular properties and cell line’s genomic characterizations to learn meaningful representations. Although deep learning can model high-dimensional biological data to capture the complex, non-linear patterns (Baptista et al. 2021, Stephenson et al. 2019), it is not well suited for omics data as deep learning has shown to fare poorly with sub-par performance on structured tabular data (Borisov et al. 2022, Shwartz-Ziv et al. 2022).

Recently, generative large language models (LLM) have exhibited unprecedented capabilities on a broad array of NLP tasks (Chang et al. 2023, Zhang et al. 2023, Liu et al. 2023). LLMs are first pretrained on large text corpora to acquire knowledge of the general language features and then fine-tuned on domain-specific datasets for downstream tasks, known as the *pretrain-and-finetune* paradigm. More recently, *pretrain, prompt and predict* has emerged as a new paradigm for the GPT (Generative Pre-trained Transformer) family of the autoregressive LLM model, which can be applied for downstream tasks using task-specific input templates, known as prompts, under the zero or few shot settings without updating the model parameters. Unlike standard deep learning models which focus on automatic feature engineering from the data for optimization on downstream tasks, GPT-based LLM models exploit input engineering that leverages textual prompts to reformulate downstream tasks through the incorporation of task-specific details.

Recent studies have noted the potential of GPT in the biomedical domain focused on information extraction (IE) and NLP tasks that entail processing unstructured biomedical texts (Gutierrez et al. 2022, Moradi et al. 2021). The applicability of GPT for DSP, however, is challenged by the structured pharmacogenomics databases, which are organized in a two-dimensional grid format as tables of rows and columns. Given the inherent heterogeneity of this tabular data with mixed column data types, adapting GPT for DSP necessitates specialized prompt design. This is a non-trivial undertaking as it involves linearizing the structured feature attributes, which could be accompanied by high-dimensional genomic values, into a natural language sentence that would elicit the most accurate drug response.

This study aims to bridge the current research gap in the application of GPT for predicting cancer drug response based on structured pharmacogenomics data. As illustrated in **Figure 1**, we utilize prompt engineering to design relevant templates that demonstrate how tabular features of drug and cell line can be integrated into GPT’s input prompt, facilitating GPT’s performance optimization for DSP through evaluation under a wide range of learning approaches. We further investigate GPT’s adaptability in various experimental settings to realize its full potential for DSP. To the best of our knowledge, this work marks the first comprehensive study of GPT’s deployment on structured pharmacogenomics data for the DSP task. The promising performance of GPT offers a new direction in advancing AI-based precision oncology through the intersection of generative LLM with structured pharmacogenomics data.

**Figure 1.**
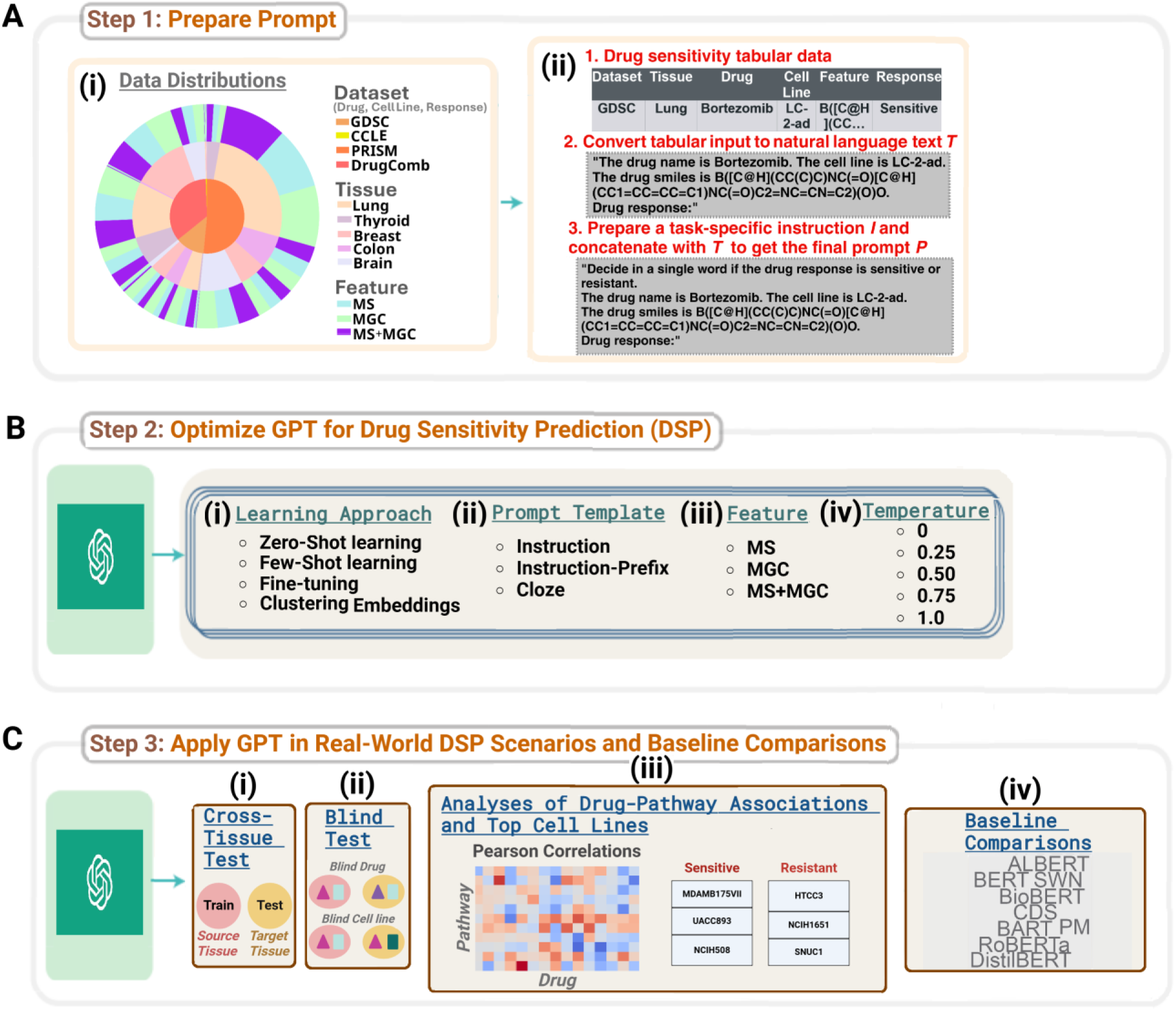
An overview of our proposed SensitiveCancerGPT framework. (A) (i) The statistics of the pharmacogenomics datasets presented as a nested pie plot. The total distribution of the cell line drug response within each dataset is shown as the innermost plot, the tissue distributions within each dataset as the middle plot and the feature distributions within each tissue cohort as the outermost plot. (ii) Workflow of prompt preparation from structured pharmacogenomics data. Using the column names and corresponding values in the tabular data, we first convert each row to a natural language text *T*. Note that the last column is left blank for the model to predict. We then prepare a task-specific instruction *I* and concatenate it with *T* to get the final prompt *P*. (B) To optimize GPT’s performance for the drug sensitivity prediction task (DSP), we evaluate on four different methodological factors: (i) learning approach, (ii) prompt template, (iii) feature and (iv) temperature. (C) We assess GPT’s generalization capability for DSP under diverse real-world experimental settings: (i) cross-tissue evaluation, (ii) blind tests, (iii) analyses of drug-pathway associations and top cell lines and (iv) baseline comparisons.

### 1.1. Statement of Significance

#### Problem

Drug sensitivity prediction (DSP) aims to predict drug response in cancer patients for personalized treatment. Large-scale pharmacogenomics datasets accumulated in high-throughput screening technologies provide a valuable resource to fulfill this task. As a result, great computational efforts have been made to analyze these data by using machine learning to build predictive models for DSP that would benefit anti-cancer therapeutics.

#### What is already known

Generative large language models (LLM) have exhibited exceptional ability in comprehending and generating unstructured textual data in the general domain, while they have been hardly applied on structured pharmacogenomics data to facilitate drug response prediction.

#### What this paper adds

This study investigates the diverse learning capabilities of the Generative Pre-trained Transformer (GPT) to analyze the genomic and molecular patterns of drug-cell lines in structured pharmacogenomics databases by introducing domain-specific novel prompt engineering to ensure effective DSP performance. We demonstrate the feasibility of GPT through comprehensive evaluations under different experimental settings including (1) cross-tissue generalization, (2) blind tests, (3) baseline comparisons and (4) analyses of drug-pathway associations and top sensitive/resistant cell lines.

## 2. Materials and methods

### 2.1. Benchmark pharmacogenomics datasets

We conduct our experiments on four publicly available pharmacogenomics resources – GDSC, CCLE, DrugComb and PRISM – which we categorize according to the respective indication of the drugs into principally two different types, oncology and non-oncology. GDSC, CCLE and DrugComb were screened against oncology drugs to test the anti-cancer sensitivity of cell lines, while PRISM screened predominantly non-oncology drugs for anti-cancer repurposing. The Genomics of Drugs Sensitivity in Cancer (GDSC) (Yang et al. 2012) comprises screening response data from two phases: GDSC1 includes 987 cancer cell lines and 320 compounds; GDSC2 includes an additional 809 cancer cell lines and 175 compounds. We use GDSC2, in which cancer cell lines are characterized by genetic features, such as the mutation state. The Cancer cell line Encyclopedia (CCLE) (Barretina et al. 2012) contains a large-scale genomic data (e.g., gene expression) for 947 human cancer cell lines and response data for around 500 of the cell lines to 24 drug compounds. The DrugComb (Zagidullin et al. 2019, Zheng et al. 2021) dataset includes data on synergy and sensitivity of drug combinations. We use the single drug sensitivity data that contains 717,684 single drug screenings from 37 studies. For each drug-drug sample in the dataset, we pair each drug with the cell line to form two separate drug-cell line inputs and label each with the corresponding single drug IC50 score. The secondary PRISM Repurposing dataset (Corsello et al. 2020) is a drug repurposing database that includes 1448 drug compounds, out of which majority were for non-oncology purposes, screened against 499 cell lines. Detailed descriptions of the datasets can be found in **Supplementary Note 2**.

#### 2.1.1. Stratification by cancer type

We stratify the drug-cell line pairs within each dataset by the tissue-of-origin to reinforce GPT’s DSP capability across a diverse spectrum of patients with different cancer types. In particular, we consider the following five tissue types which were available across all four datasets: lung, thyroid, breast, brain, colon/stomach. The data distributions are shown in **Figure 1A(i)** (**Supplementary Table 1**) and further details are provided in **Supplementary Note 3**.

### 2.2. Generative LLM used

Large Language Models (LLM) refer to scaled-up Transformer-based pretrained language models. The Generative Pre-trained Transformer (GPT) variants from OpenAI are decoder-based autoregressive family of LLMs and includes, at the time of this study, GPT-3, GPT-3.5 and GPT-4. These models are accessible through OpenAI API at a usage fees. Although the successor LLMs of GPT-3 are known to demonstrate stronger capabilities, they incur higher inference cost. To balance the cost-effectiveness and DSP performance of GPT, we carried out an empirical analysis (**Supplementary Note 4**) comparing evaluations between GPT-3, GPT-3.5 and GPT-4. We found that GPT-3.5 and GPT-4 performed the same as GPT-3 with no additional improvements. Hence, we opt for GPT-3 in all our experiments.

### 2.3. Learning approach

We employ four different paradigms to adapt GPT-3 for downstream DSP analysis: (i) Fine-tuning trains and then tests the pretrained GPT-3 model on prompt-completion pairs, (ii) zero-shot learning directly runs the pretrained GPT-3 model on textual prompt inputs from the test set to generate the drug sensitivity response during inference, (iii) few-shot is akin to zero shot but additionally integrates few examples into the prompt and (iv) clustering embeddings refines the GPT-3-derived textual embeddings by fitting a Bayesian Gaussian Mixture Model (BGMM) (Bishop C 2006). Please refer to **Supplementary Note 5** for detailed descriptions.

### 2.4. Prompt engineering

Prompting GPT-3 for DSP refers to providing the relevant pharmacogenomics knowledge via carefully designed textual templates to guide the model’s response generation. The prompt preparation steps are illustrated in **Figure 1A(ii)**. The pharmacogenomics data is formatted as a table of M rows of drug-cell line instances and N columns. Note that the first row defines the headers corresponding to the column names. We employ relevant prompt engineering of the tabular pharmacogenomics data specific to each learning approach.

#### 2.4.1. Zero-shot

A standard prompt *P* could be crafted with the following elements: task-specific instruction *I*, input *T*, context *C* and output placeholder *R*. We perform table serialization (Hegselmann et al. 2023) by converting each row in the test set to a natural language text *T* based on the headers and the corresponding column values. Note that the last column’s value is considered as *R* and left blank. We then prepare *I* and concatenate it with *T* to get the final prompt *P*, which is directly inputted into GPT-3 to generate the drug response.

#### 2.4.2. Few-shot

We expand the zero-shot prompt template *P* with context *C*. The context *C* constitutes *k* pairs of drug-cell line and ground truth response examples. We experiment with three different prompt design templates. We devise the three templates based on the following: inclusion of *I*, format of *T* and *C*, and order of *R*. As such, *instruction prompt* includes *I*, has a sentence format for *T* and *C*, and places *R* at the end: *P = I + T + C + R*. Here, the *+* sign denotes concatenation followed by a new line. The *instruction-*p*refix prompt* is similar to instruction prompt, however, *T* and *C* have a concise format as we prepend the headers to the respective column values with colon prefix: *P = I + T’ + C’ + R*, where *’* indicates the concise format. Lastly, *cloze prompt* does not include *I*, inserts *R* within *T* (analogous to filling in the blank), and maintains sentence format for *T* and *C*: *P = TR + C*. We illustrate each prompt template with an example in **Supplementary Figure 2**.

#### 2.4.3. Fine-tuning

We prepare the training and test data in the form of prompt-completion pairs. The prompt is represented using a concise prompt template similar to instruction-prefix and the completion is the ground truth sensitive/resistant label.

### 2.5. Prompt features

The content within *T* and *C* could be grounded with pharmacogenomics knowledge derived from the ‘Feature’ column. We assess the informativeness of two types of features - *molecular structure information* (*MS*) and *additional molecular or genomic context* (*MGC*) - that are integrated into the prompt template with the drug-cell line input (*Basic Information* (*BI*)). This leads to three feature groups in total (BI + MS, BI + MGC, BI + MS + MGC), that are evaluated per dataset. The feature descriptions can be found in **Supplementary Note 6** and the feature-ablated dataset sizes are shown in **Supplementary Table 1**.

## 3. Experiment Setup

### 3.1 Task Formulation

We formulate drug sensitivity prediction as a binary classification problem to predict if a drug-cell line pair is *sensitive* or *resistant*. For the drug-cell line pairs in each dataset, we adopt the corresponding IC50 (the half maximal inhibitory concentration) values to gauge the drug sensitivity. To convert the IC50 drug response values to binary labels, we adopt the strategy employed in (Chang et al. 2018) and set a fixed threshold θ = −2, where ln(IC50) < θ is considered sensitive, and resistant otherwise.

### 3.2. Evaluations

#### 3.2.1. Optimization of GPT’s DSP performance

We investigate the influence of four methodological factors that shape the DSP performance. These include learning approach, prompt template, feature and temperature. For learning approach, prompt template and feature, we use the settings outlined in **Sections 2.3**, **2.4.2** and **2.5**, respectively. For GPT-3’s temperature setting, we evaluate on values in increment of 0.25 between 0 and 1. Analysis is carried out separately on each factor by evaluating GPT-3 relative to the settings associated with that factor, while the other three factors’ settings are kept fixed: fine-tuning for learning approach, instruction-prefix for prompt template, MS for feature, and 0 for temperature. Only in the assessment of prompt template, we set learning approach to few-shot.

To adequately probe how the prompt context *k* contributes to DSP performance, we further conduct subanalyses for few-shot learning to (1) examine the impact of increasing *k* by varying as 1, 5, 10 or 15 examples, and (2) compare the selection strategy of *k*.

#### 3.2.2. Application of GPT in Real-world DSP scenarios

##### 3.2.2.1. Cross-tissue generalization

In the within-tissue analysis, we fine-tune and test drug-cell line pairs originating from the same tissue type, assuming that the data distributions of the training and test sets are the same. However, for certain tissues (e.g., rare cancer), the cold-start problem is inevitable where cell lines are scarce and thus cannot suffice both the training and test sets. In such scenarios, it would be beneficial if the knowledge acquired during fine-tuning on a common tissue dataset could generalize to a rare tissue dataset. To mimic this real clinical setting where training and test samples are available for different tissues, in the cross-tissue analysis per dataset, we fine-tune the model using the largest tissue cohort in that dataset and subsequently apply the fine-tuned model to predict drug response on the remaining four tissue cohorts.

##### 3.2.2.2. Blind tests

In the standard experiment setup, the drug-cell line pairs are randomly split into training and test sets, so it is possible for the same drug or cell line to exist in both the training and test sets. This enables GPT-3 to predict the sensitivity of a new cell line to a drug, having already known the same drug’s effect on a different cell line during training, and vice versa. We conduct stringent splitting setups by constraining the drug or cell line from existing simultaneously in both the training and test sets. That is, (i) in the *blind test for cell line*, we predict the sensitivity of a new test cell line that is unseen in the training set to a test drug that is seen in the training set and (ii) in the *blind test for drug*, we predict the sensitivity of a test cell line that is seen in the training set to a new test drug that is unseen in the training set.

##### 3.2.2.3. Analysis of the drug-pathway associations

When a drug exerts its effects on a cell line, it is known to affect a related pathway rather than a single target (Wang et al. 2021). Henceforth, we leverage GPT-3’s pretrained text embeddings to generate the heatmap of Pearson correlations between the predicted drug responses (IC50) and pathway activity scores, with the possibility to obtain biological insights toward the drug response mechanism. More details are provided in **Supplementary Note 7**.

##### 3.2.2.4. Analysis of the top cell lines

In this case study, we want to investigate if GPT-3’s confidence of the drug response predictions (log probabilities) aligns with the reported experimental evidence in external databases. For illustrative purpose, we consider the cell lines of the drug Afatinib in the GDSC dataset as the test drug-cell line samples and evaluate them using the fine-tuned GPT-3 model. The log probabilities outputted by GPT-3 are sorted to find the cell lines most sensitive and resistant to Afatinib. To verify if the top cell lines predicted by GPT-3 are corroborated by relevant evidence, we use the online analysis portal (Fekete et al. 2022) which provides ranking of cell line models across four drug screening datasets. We consider Afatinib as it was evaluated in our benchmark dataset, GDSC, and was further validated in (Fekete et al. 2022).

#### 3.2.3 Baseline Comparisons

For baseline comparisons, we consider the following Transformer-based pretrained language models: *BERT* (Devlin et al. 2018), *BART* (Lewis et al. 2019), *BioBERT* (Lee et al. 2020), *DistilBERT* (Sanh et al. 2019), *RoBERTa* (Liu et al. 2019) and *ALBERT* (Lan et al. 2019). We also compare against three existing drug response models: *SWNet* (SWN) (Zuo et al. 2021), *PaccMann* (PM) (Manica et al. 2019) and *ConsDeepSignaling* (CDS) (Zhang et al. 2021). Please refer to **Supplementary Note 8** for baseline descriptions and implementation details.

## 4. Results

**Figure 2A** shows the evaluation results of the methodological factors over different datasets and tissues as box-plots. For ease of comparison among the settings within each factor, we consider the mean F1, which is computed by averaging the results pertaining to a setting across all the datasets and tissues. In the comparison of different learning approaches in Figure 2A**(i)** (detailed results in **Supplementary Figure 4**), fine-tuning attains the best performance with a mean F1 of 0.84 and provides significant improvements over zero-shot (0.24, p-value= 2.3e-18) and few-shot (0.66, p-value=1.5e-10), while rivaling with competitiveness against clustering (0.83, p-value=0.77). A comparison of the different prompt templates evaluated in the few-shot setting is presented in **Figure 2A(ii)** (detailed results in **Supplementary Figure 5**). We observe the best results with instruction-prefix (mean F1 0.68), resulting in performance gap of 4% against instruction prompt (mean F1 0.65), albeit a not statistically significant difference (p-value=0.40), and a more pronounced gap of 22% (p-value= 0.02) against cloze prompt (mean F1 0.55). We analyze the contribution of different combinations of features to the DSP performance in **Figure 2A(iii)** (detailed results in **Supplementary Figure 6**). We use performance with BI as a reference point to evaluate the change in performance instigated by the three feature sets. Overall, incorporating the drug’s molecular representation in the SMILES notation (i.e., BI + MS) does not provide any added predictive power, but rather causes a 3% decrease (p-value=0.46) of the mean F1 performance from 0.84 to 0.81. With regard to the BI + MGC feature, prompt context derived from drug’s synergy information in DrugComb dataset and drug’s mechanism of action in PRISM lead to performance boosts (13.4% mean F1 increase, p-value=0.02 and 2.3% mean F1 increase, p-value=0.28, respectively). Gene mutation information in GDSC does not cause any drastic change in performance and, conversely, gene expression in the CCLE dataset has a negative effect (3.5% mean F1 decrease; p-value=0.31). Lastly, contextualizing the prompt with both MGC and SMILES (i.e., BI + MS + MGC) generally lags behind in mean F1 performance in comparison to the absence of SMILES (BI + MGC) by 1% (p-value=0.72). In the temperature analysis (**Figure 2A(iv)**), the performance remains unaffected across the different settings. Based on the best evaluation results of the methodological factors, we test GPT-3’s DSP capability under the optimal methodological settings on the five tissues cohorts across the datasets. The F1 and per-class F1 results in **Figure 3** suggest that GPT-3 performed competitively on the GDSC and PRISM datasets across all the tissues, while the worst on the CCLE dataset with performance declines in F1 (p-value=0.15), F1-Sensitive (p-value=0.02) and F1-Resistant (p-value=0.24).

**Figure 2:**
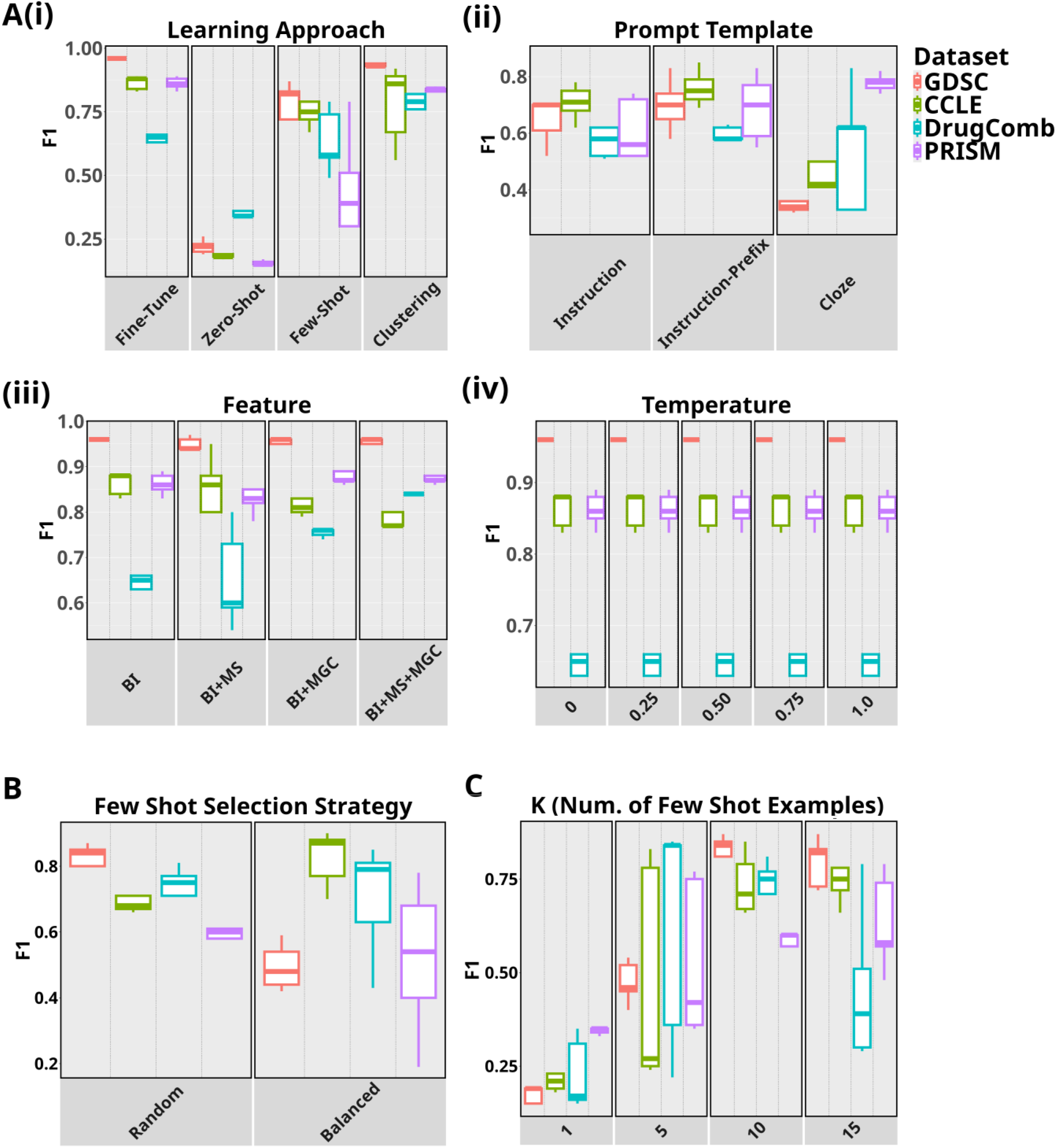
(A) Performance of GPT under different settings associated with the following methodological factors: (i) learning approach, (ii) prompt template, (iii) context and (iv) temperature. (B) Few-shot performance comparisons of datasets on varying the number of demonstrations, *k*, in increments of five from 1 to 15. (C) Performance comparisons between two different selection strategies under the few-shot setting.

**Figure 3:**
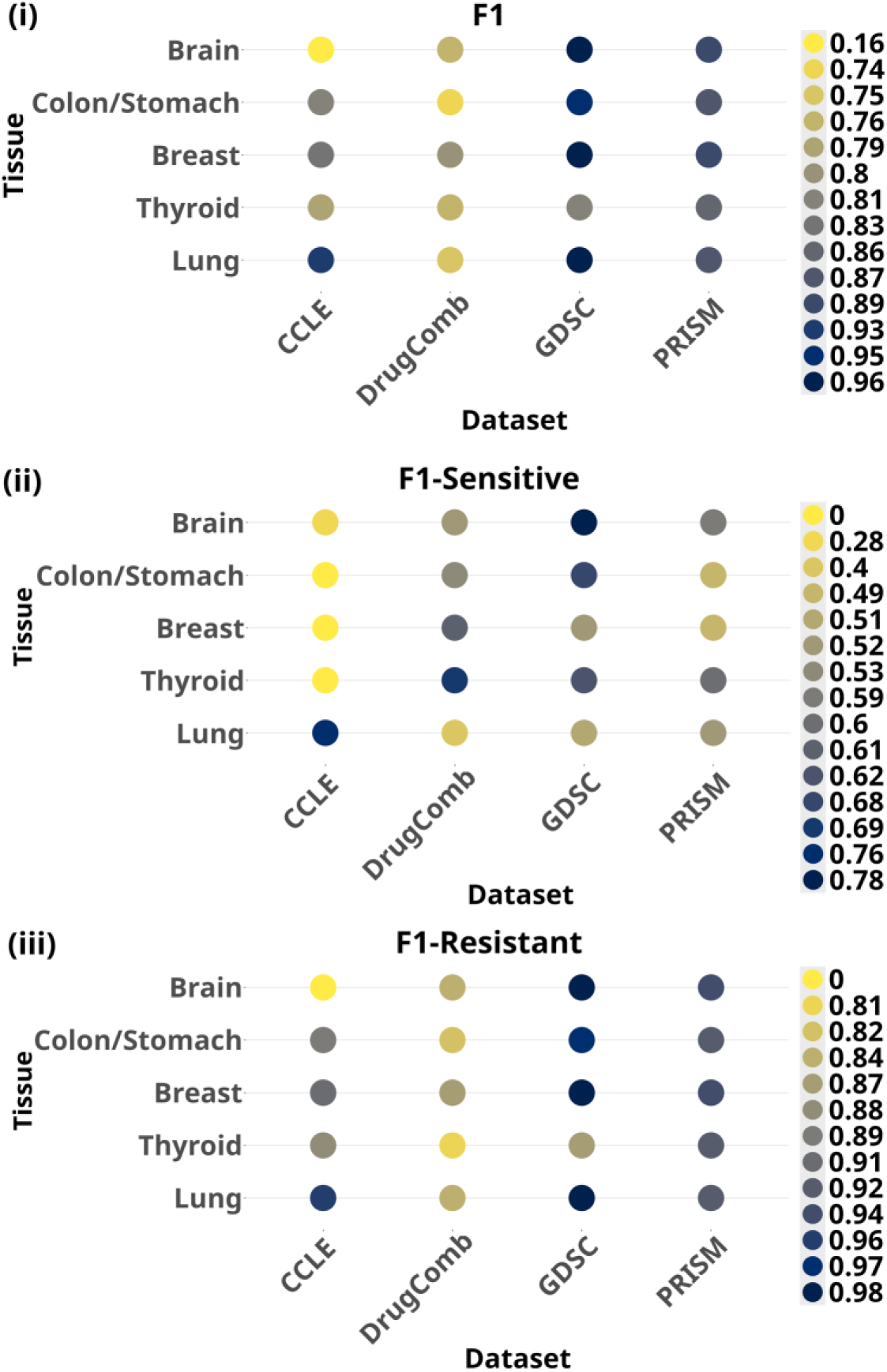
Performance of GPT with optimal settings on five tissue cohorts across different pharmacogenomics datasets evaluated with (i) F1 (ii) F1-Sensitive and (iii) F1-Resistant. F1 is the micro-averaged F1 score and F1-Sensitive and F1-Resistant are the F1 scores for the positive and negative classes, respectively. We used scikit-learn (Pedregosa et al. 2011) for the computation of evaluation metrics.

Comparison of performances with different number of demonstration examples *k* in **Figure 2B** reveals that GPT-3’s in-context generalizability is sensitive to *k* and prompting with more drug-cell line examples (i.e., higher *k*), in general, is beneficial. The experiments comparing between the random and balanced selection strategies of *k* in **Figure 2C** provide mixed results across datasets. Balanced selection strategy surpasses random selection in CCLE and PRISM, while the reverse is true for GDSC and DrugComb, although neither of them is statistically significant.

We compare GPT-3’s within-tissue and cross-tissue prediction abilities across the tissues within each dataset as violin plots in **Figure 4A**. In GDSC and PRISM evaluations, the cross-tissue performances are able to uphold comparable results to the corresponding evaluations in the within-tissue setting. Whereas evaluations on the CCLE and DrugComb datasets attain heightened cross-tissue performances with significant mean F1 performance gains of 8.1% (p-value=0.001) and 18.7% (p-value=0.009), respectively.

**Figure 4:**
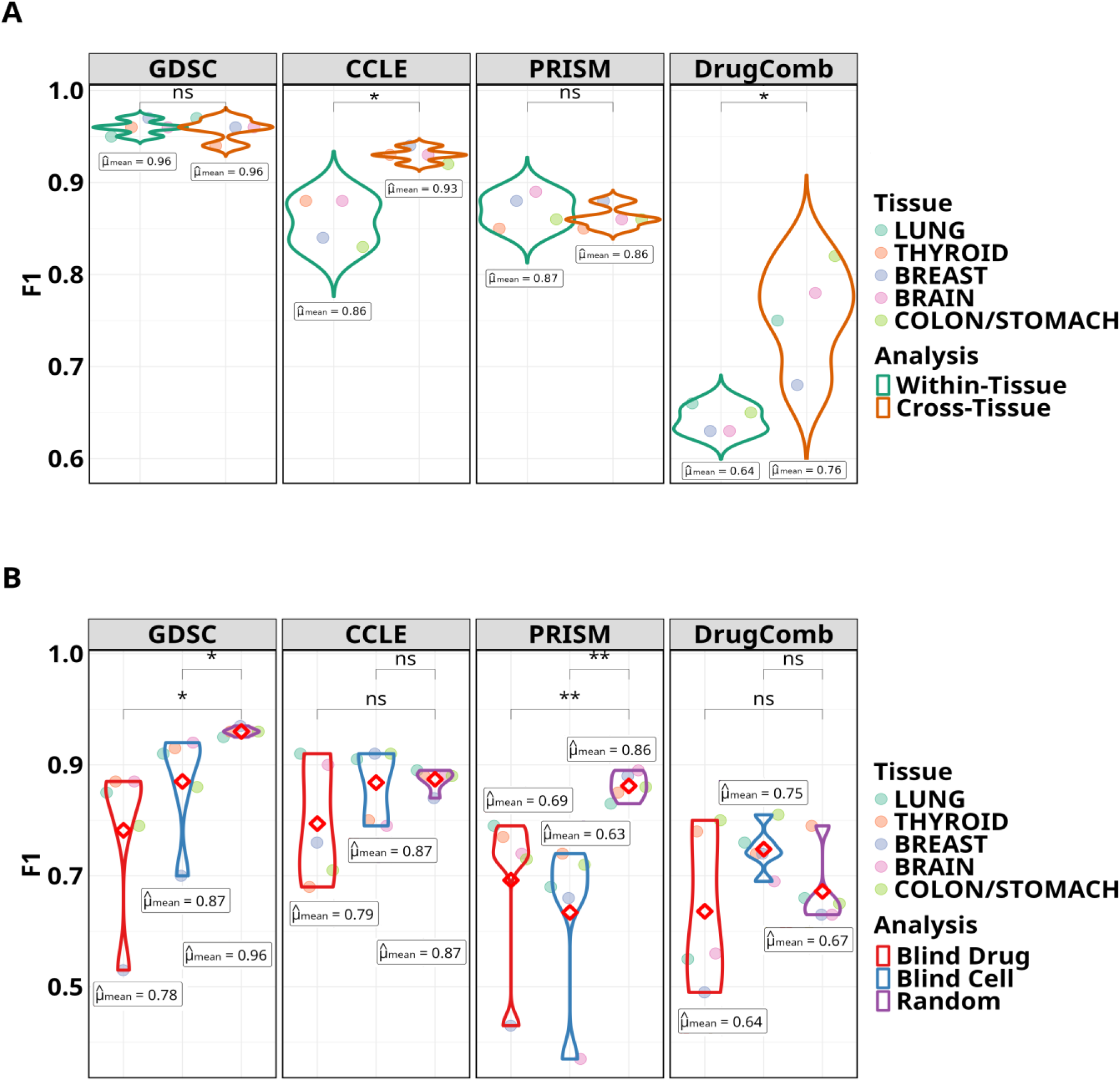
Performance comparisons in the form of violin plots for the (A) analysis between within-tissue and cross-tissue evaluation settings and (B) blind test analyses for drugs and cell lines. In each type of analysis, GPT is evaluated on the five tissue cohorts associated with each dataset separately. So, the annotated mean (also shown as the red diamond shape) is computed by averaging the results across all tissues within a dataset. The degree of statistical significance is indicated by the number of asterisks and a not statistically significant difference by ‘ns’.

The violin plots in **Figure 4B** depict GPT-3’s blind test generalizations (Blind Drug, Blind Cell) over the five tissue cohorts within each dataset. The baseline performance was established by evaluation on the random test split (Random). GPT-3 was able to attain competitive performances (as not statistically significant difference) on par with the corresponding random evaluations for CCLE and DrugComb. In contrast, the blind evaluations underperform on PRISM (p-value=0.01) and GDSC (p-value=0.05). Comparing between blind drug and blind cell evaluations, generally, GPT-3 lags behind in accurately predicting response of drug-cell line pairs that include new drugs, with mean F1 performance gains toward blind cell tests for the following datasets: GDSC (11.53%; p-value=0.29), CCLE (10.12%; p-value=0.23) and DrugComb (17.1%; p-value=0.13).

In **Figure 5A** (detailed results in **Supplementary Figures 7 and 8**), we compare the F1 and per-class performances of GPT-3 with baseline models. GPT-3 outperforms baselines with mean F1 gains of 5.4% in GDSC, 13.6% in CCLE, 5.3% in DrugComb and 5.3% in PRISM (p-values reported in **Supplementary Note 9**). Additionally, GPT-3 was able to maintain superior F1-Sensitive performance against BERT (p-value=3.6e-05), BART (p-value=0.05), BioBERT (p-value=0.00026), DistilBERT (p-value=0.15), ALBERT (p-value=0.13), RoBERTa (p-value=0.03), CDS (p-value=0.97) and PM (p-value=0.012). Although SWN’s F1 and F1-Sensitive scores surpass GPT-3’s, the differences are not statistically significant (p-value=0.06 and p-value=0.59 respectively). In **Figure 5B**, we examine the F1-Sensitive performances of GPT and baselines with varying positive class distributions. In a label scarce setting with only 20% of sensitive drug-cell lines available in GDSC’s tissue cohorts, GPT-3 is seen to perform significantly more robustly compared to the baselines (p-value=1.1e-22).

**Figure 5:**
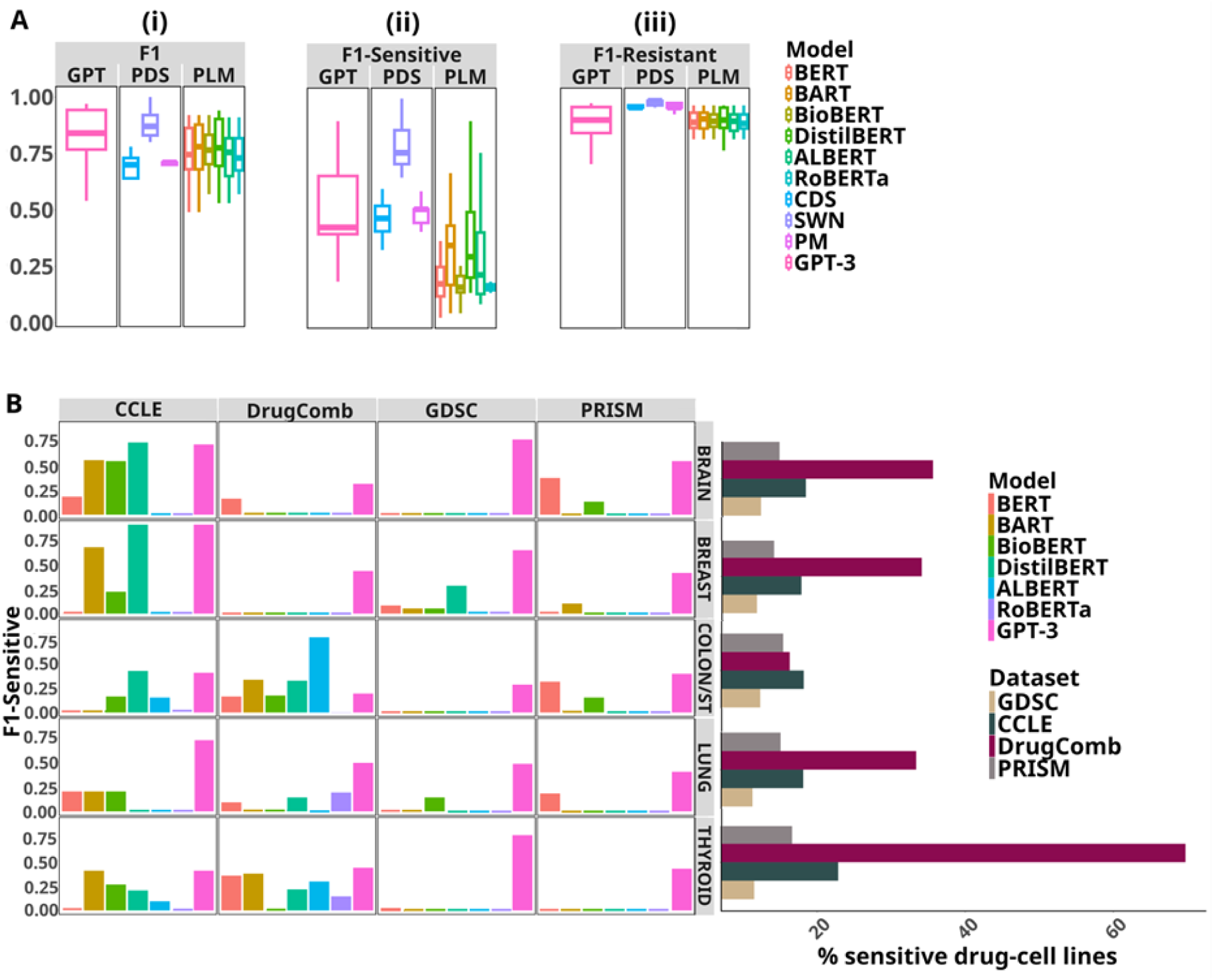
(A) Performance comparisons of GPT with two types of baseline models, previous drug response prediction models (PDS) and pretrained language models (PLM). GPT and the PLM baselines are fine-tuned. The F1 and per-class F1 reported are averaged across all five tissues’ evaluations across the four datasets for each model. Detailed results are available in Supplementary Figure 7 and Supplementary Figure 8. (B) F1-Sensitive performances of GPT and baselines with varying positive class distributions.

We visualize heatmap of the Pearson correlations between the predicted drug responses (IC50) and pathway activity scores of the cell lines generated by PROGENy (Schubert et al. 2018) in **Figure 6A**. The computed p-values assert high similarity between the corresponding correlation distributions (i.e., predicted (**Figure 6A(i))**, actual (**Figure 6A(ii))**) across all pathways, thus justifying that GPT-3-derived text embeddings could facilitate in both the accurate prediction and explainability of the drug response. Importantly, we observed that the drugs with relatively low IC50 scores were sensitive to major pathways and these findings align with those in the previous research (**Section 5**). In a case study analysis, we further corroborate GPT-3’s generalization in terms of its predicted log probabilities for the drug Afatinib by ranking its top cell lines in **Supplementary Table 3**. GPT-3 was able to predict drug response with high confidence for the most sensitive and resistant cell lines to the drug Afatinib, which have been validated in previous experimental studies (Fekete et al. 2022).

**Figure 6:**
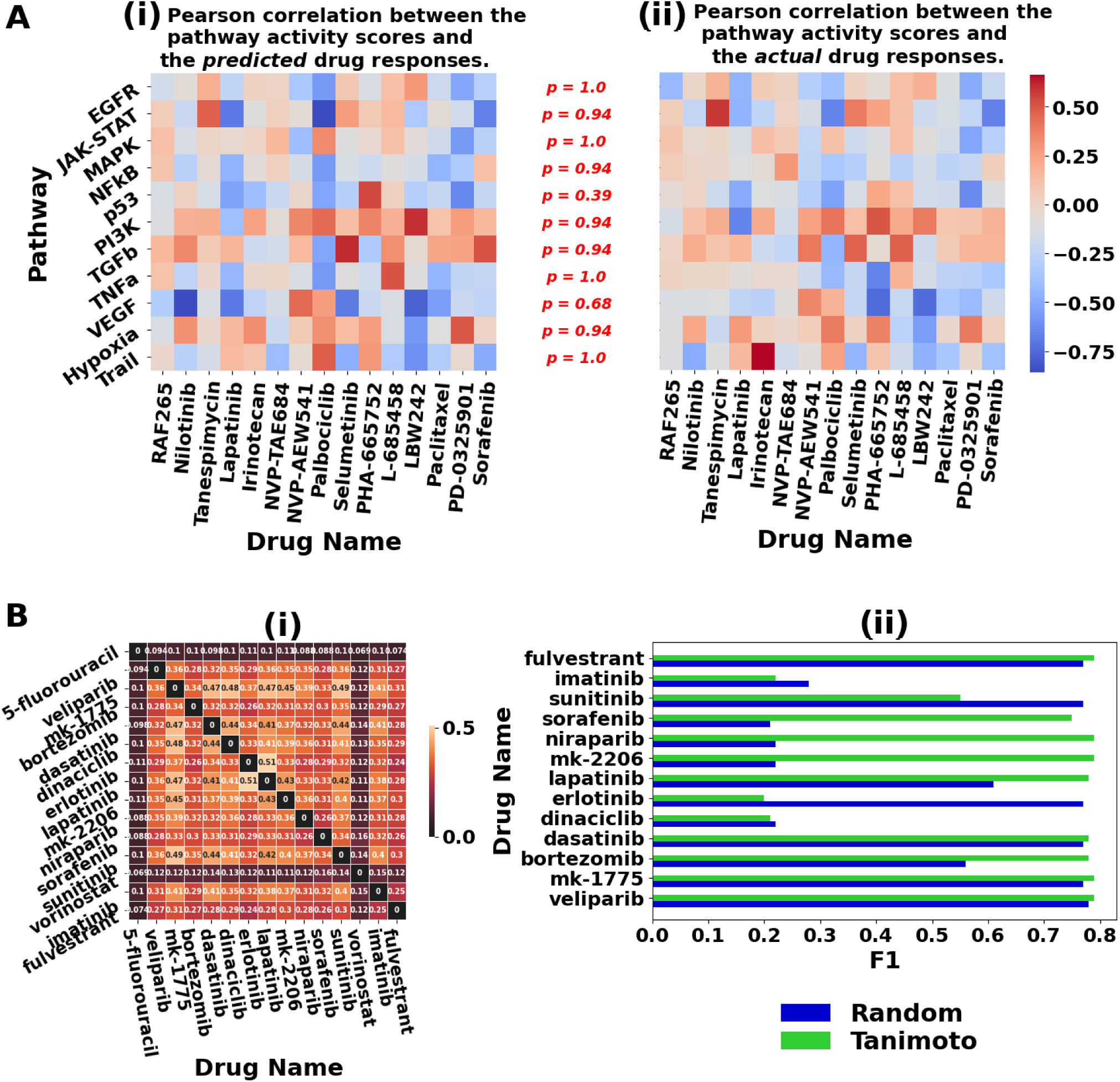
(A) Drug-pathway associations in the CCLE dataset. Negative (blue) and positive (red) correlations correspond to ‘assistant’ and ‘resistant’ associations, respectively. The p-values, annotated in red (i.e., p =), are computed using the Kolmogorov-Smirnov test and indicate distribution similarity between the predicted (subplot (ii)) and actual (subplot (ii)) drug-pathway associations. Note that the null hypothesis (p > 0.05) is that the two distributions are similar. (B (i)) Heatmap visualization of Tanimoto correlations between drugs. (ii) Few-shot classification performance using Tanimoto-based (i.e., structurally similar drugs) selection for in-context examples in the prompt.

## 5. Conclusion and Discussion

The advent of LLMs have revolutionized the field of natural language processing. However, their applicability to biomedical tasks with structured data is still missing or limited in the current research. This work showcases the feasibility and efficacy of a generative LLM, namely GPT-3, in the prediction of cancer cell line’s response to oncology and non-oncology drugs, with a focus on leveraging structured pharmacogenomics data. To account for the crucial role prompt engineering plays in LLM’s performance, this study provides a nuanced exploration of methodological factors in tailoring prompt engineering to optimize the DSP task. Additionally, it contributes valuable insights toward GPT-3’s application in DSP that cater to diverse real-world evaluation scenarios, paving the way for personalized treatment.

We investigated the effect of the type of feature integrated into the prompt context on GPT-3’s performance. The results suggest that drug descriptor in the form of SMILES can be, in general, detrimental from GPT-3’s generalization perspective. The SMILES notation linearly represents the atom connectivity in a chemical compound as a text sequence. However, unlike a natural language text, the characters in a SMILES sequence correspond to topological characteristics, such as ring-closure or branches, that cannot be treated as isolated entities. Furthermore, the presence of repetitive tokens in the SMILES string causes the atoms of a molecule to be indistinguishable in the token space (Ucak et al. 2023). We reason that the drop in performance was as a result of GPT-3’s failure in extracting such nuanced features from the SMILES-incorporated prompt. This is likely attributed to GPT-3’s base vocabulary not accounting for SMILES grammar, that hinders the frequency-based Byte Pair Encoding tokenization (Sennrich et al. 2015) from effectively splitting the SMILES sequence into meaningful constituent elements without losing the molecule’s chemical validity. Unsupervised pretraining of GPT-3 using a large SMILES corpus could possibly warrant performance enhancement. In contrast, supplementing the drug-cell line input with drug’s functional feature aids in performance improvements. In relation to DrugComb, drug synergy has shown to have a positive correlation with drug sensitivity such that cell lines involved in synergistic drug combination are also significantly more sensitive to the drugs (Narayan et al. 2020). Similarly, in PRISM evaluation, drug sensitivity is affected by the drug’s MOA as it identifies the binding target and whether the therapeutic effects will be enabled by the activationor inhibition of the target’s function. Although genomic mutations are deemed to provide significantly discriminative patterns across cell lines (Yang et al. 2012), it only benefitted the DSP performance on GDSC’s lung cohort evaluation. The use of gene expression feature was dominated by a decline in CCLE’s performances across tissues. We rationalize CCLE’s drop in performance to the “small n, large p” problem (Nguyen et al. 2021) wherein given the smaller number of drug-cell line pairs in the dataset, the gene expression sequence increases the overall feature dimensionality (i.e., prompt context length) causing obstruction in the model’s learning and resulting in possible overfitting.

We examined GPT-3’s DSP capability through a diverse spectrum of learning approaches. Zero-shot learning is a training-free approach that generates the drug response by conditioning on the input prompt and relying on the knowledge already encoded in GPT-3’s parameters. The unfavorable zero-shot generalization of GPT-3 suggests that it was not exposed to sufficient pharmacogenomics-related content during its pretraining phase. However, augmenting the prompt with a few drug-cell line examples via few-shot prompting, that demonstrate the input-output mapping existing in the drug sensitivity datasets, initiates relevant learning and hence is seen to elevate the overall performances. This is consistent with the findings in the general domain where GPT-3 excelled in NLP tasks in the few-shot setting (Brown et al. 2020). Our analysis of prompt engineering in the few-shot setting suggests that providing explicit instruction and conforming to a concise in-context exemplar format (i.e., instruction-prefix prompt) elicits the most accurate drug response prediction from GPT-3. Drugs with similar chemical structure have shown to exhibit similar sensitivity patterns (Zhang et al. 2015). We emulate this finding in GPT-3’s in-context learning (**Supplementary Note 10**) and observe that demonstration examples selected based on structurally similar drugs enhance GPT-3’s few-shot classification, in general (**Figure 6B**). Fine-tuning entails training GPT-3 on supervised pharmacogenomics datasets. The domain-specific knowledge acquired thereafter visibly causes substantial spike in the performances of all datasets. In the weakly supervised paradigm, we iteratively refine GPT-3’s pretrained embeddings by clustering via BGMM. This benefits GPT-3’s DSP generalization in comparison to inference contingent on its pre-existing knowledge (zero-shot), resulting in significant performance gains (mean F1 increase from 0.24 to 0.83). More importantly, given that the pseudo-labels for cluster initialization are derived from fine-tuning, their mean F1 performances are comparable and at times clustering even exceeds fine-tuning performance, exemplified in the following evaluations: DrugComb’s lung, thyroid, breast, brain and CCLE’s breast.

GPT-3 has demonstrated remarkable out-of-distribution generalization, underscoring its ability to predict drug response for cancer types completely unobserved during training. The cross-tissue evaluation corresponds to simulating the challenging cold-start scenario in precision oncology, where a large training dataset is not readily available for certain tissue types but only a few samples could be exploited for testing. Harnessing transfer learning that relies on fine-tuning GPT-3 on a large amount of common tissue drug response data has potential use case in recommending treatments to patients with rare cancers. We further evaluated GPT-3 in challenging data splitting strategies. The blind cell test aims to find candidate drugs among all the drugs the model is trained on for a new cell line (i.e., new patient) and has direct utility in personalized cancer treatment. The blind drug test, on the other hand, is designed to generalize for a new drug to known cell lines and could be useful in repurposing non-cancer drugs for cancer treatment, as well as novel drug discovery (Partin et al. 2023). Our empirical results suggest that GPT-3’s prediction performances under the stricter blind partitioning settings are comparable to those with the baseline random splitting on CCLE and DrugComb. We further observed that GPT-3’s generalization in the blind cell analysis is generally superior to the blind drug test, although the difference is not statistically significant. Even though blind cell analysis ensures that the test set does not contain any cell lines already present in the training set, it is possible to share biological characteristics if the unknown test cell lines and known train cell lines happen to belong to the same tissue. On the contrary, the relatively large chemical space of drug compounds makes it difficult for GPT-3 to learn generalizable features for an unknown drug.

We benchmarked GPT-3 against a range of established baseline models for a rigorous performance assessment on the drug sensitivity prediction task. GPT-3 outperformed the baselines by achieving a higher mean overall F1 score, with mostly statistically significant performance differences for evaluations on the GDSC, CCLE and PRISM datasets. Importantly, F1-Sensitive is the most relevant metric for drug response prediction as it shows the correctly classified sensitive drug-cell line pairs, knowledge of which could accelerate treatment optimization and drug repurposing. On inspecting the Transformer-based language model baseline generalizations for the sensitive class, there are some instances where GPT-3’s overall F1 score is below the baseline performances (e.g., thyroid assessment in CCLE, brain assessment in DrugComb), however, it is procured at the expense of relatively lower corresponding F1-Sensitive scores, in general. This indicates that GPT-3 is not only able to successfully distinguish the sensitive samples from the resistant ones, but also strikes a good balance in accurately identifying drug-cell line pairs belonging to the minority sensitive class. This could be attributed to GPT-3’s larger size (175B parameters) and the fact that it has been pretrained on much more data (300B tokens), that helps to encapsulate domain-specific signals better during fine-tuning. In comparison to the existing drug response models (i.e., CDS, SWN, PM), GPT-3’s superior performance implicates the predictive benefit of prompt-based input engineering over the traditional feature engineering in providing task-specific context. The superimposition of sensitive class distributions of tissues across the datasets with F1-Sensitive performances (**Figure 5B**) further indicates that GPT’3’s fine-tuning capability for DSP is not constrained by label scarcity as performances on thyroid, breast, brain and colon/stomach tissues of GDSC, which has the fewest sensitive samples, supersede counterpart cohorts on DrugComb with the largest positive class distributions.

In order to accommodate GPT-3 for interpretability study, we conducted a post-hoc analysis based on drug-pathway associations through leveraging GPT-3’s pretrained text embeddings of input prompts. As a drug is known to exert its effects by influencing the gene expression in the related pathway (Wang et al. 2021), the drug-pathway associations facilitate in providing biological justifications for the predicted drug responses. We observed several instances of drug-pathway associations for which our calculated Pearson correlations are consistent with the findings in previous research. Sorafenib is reported to inhibit the tumor cell growth by blocking the MAPK pathway, indicating an assistant association (Gauthier et al. 2013). Sorafenib, a multi-kinase inhibitor, and RAF265, a RAF inhibitor drug (McCubrey et al. 2007), are found to be negatively correlated (assistant association) with the VEGF pathway (Liu et al. 2006). Analogously, Nilotinib, a tyrosine kinase inhibitor (TKI), is known to alter VEGF-induced angiogenesis pathway (Zibrova et al. 2024). We observed negative correlation between the EGFR pathway and Lapatinib, which is an EGFR inhibitor (Montemurro et al. 2007). The drug LBW242 is known to sensitize cancer cells to the antitumor effects of TRAIL (Philemon et al. 2015), indicating an assistant association.

Despite the promising performances, the safety-critical nature of the domain necessitates that some limitations need to be addressed before a generative LLM like GPT-3 can be deployed and integrated into the medical workflow. In the present study, we tested GPT-3’s interpretability by leveraging its pretrained text embeddings. A possible future direction would be to directly prompt GPT-3 via the few-shot chain-of-thought strategy (Wei et al. 2022) with the possibility to generate the reasoning chains justifying its predicted response, while incorporating human-in-the-loop to correct any reasoning errors or hallucinations. Considering the closed-source nature of GPT-3 family models’ accessibility, we anticipate future efforts of applying the systematic methodology presented in this work to investigate open-source LLMs (e.g., LLaMA) for a broader understanding of DSP evaluation.

## Supplementary Material

### Supplementary Note 1: Background

#### Computational Drug Sensitivity Prediction

The various methods employed to predict drug response in silico can be categorized into network-based approaches (Guan et al. 2019, Cheng et al. 2019) and machine learning approaches (Dong et al. 2015, Menden et al. 2013). Network-based approaches model the relationships between the heterogeneous entities in the pharmacogenomic data (e.g., drug, target, cell line) by constructing similarity or interaction graphs. Machine learning approaches, on the other hand, extract informative features from the pharmacogenomics data, based on which training and prediction of drug sensitivity is carried out. Recently, deep learning has rapidly made strides owing to its ability to automatically learn the complex, non-linear patterns from large amounts of pharmacological and cell line omics data (Baptista et al. 2021, Stephenson et al. 2019, G et al. 2020). Importantly, effective representations are learned by encoding chemical and genomic properties that could facilitate the automatic extraction of latent features informative in the drug response prediction. DeepDSC (Li et al. 2019) first learns compressed representation of gene expression data through the application of a stacked autoencoder, which is then integrated with drug chemical features in the form of molecular fingerprints to predict drug response using a feed forward neural network. DeepDR (Chiu et al. 2019) employs separate autoencoders pretrained on pan-cancer data to transform gene mutation and expression features into low dimensional representations and links them for drug response prediction. Unlike autoencoder-based approach, end-to-end drug response models have also been proposed that jointly encode and predict as a unified model. SWNet (Zuo et al. 2021) adopts a dual convergence model architecture to simultaneously encode the drug and genomic features using graph neural network and convolutional neural network, respectively. It then exploits late integration between the two representations before passing through the prediction subnetwork. Manica et al. 2019 proposes a multimodal architecture and explores various models for encoding drug representation from SMILES, while using an attention-based gene encoder.

This work explores the application of a generative LLM (i.e., GPT) in drug sensitivity prediction (DSP). Unlike standard deep learning models which focus on automatic feature engineering from the data for optimization on downstream tasks, GPT-based LLMs exploit input engineering that leverages textual prompts to reformulate downstream tasks through the incorporation of task-specific details.

#### Generative LLM for biomedical tasks

The proliferation of biomedical corpora and the idiosyncrasies of biomedical text have led to domain-specific research of LLM in biomedicine. Gutierrez et al. 2022 explored the in-context learning capability of GPT-3 for named entity recognition and relation extraction. Moradi et al. 2021 also evaluated GPT-3 in the few-shot setting but across a broader range of biomedical NLP tasks encompassing textual inference, relation classification, semantic similarity estimation, question answering and text classification. Labrak et al. 2023 performed a comprehensive assessment of four instruction-tuned LLMs (ChatGPT, Flan-T5 UL2, TkInstruct, and Alpaca) in the zero-shot and few-shot settings on 13 biomedical tasks covering diverse NLP problems: classification, question answering, relation extraction, natural language inference and named entity recognition.

However, these studies focus on processing NLP datasets that include unstructured text and their applicability to biomedical tasks with structured data is still missing or limited in the current research. Given that structured pharmacogenomics data have special data formats that a generative LLM is not naturally familiar with and cannot well comprehend as it is pretrained on textual Web data, specialized prompt design that involves linearizing the structured input into sentence to elicit the most accurate drug response is non-trivial. This work aims to bridge this research gap by introducing prompt engineering that is tailored to structured pharmacogenomics data to facilitate DSP.

#### Deep neural networks for tabular data

Tabular data is ubiquitous in many real-world domains, including healthcare (Bhatt et al. 2020, Jiang et al. 2024). Data within a table is laid out in rows and column format, with each row corresponding to an input sample and each column representing a feature attribute. Thus, accurate prediction over tabular data necessitates semantic structure understanding capabilities. The wide application of deep neural networks to tabular data is attributed to its ability in automatically learning non-linear feature representations from table cells. Several variants of the vanilla multilayer perceptron (MLP) have demonstrated improved performance on tabular data. Work by Kadra et al. 2021 applied several regularization techniques (i.e., regularization cocktail) to the MLP, a parameter-efficient ensemble of MLP was proposed in Gorishniy et al. 2024, and an enhanced MLP obtained through better default parameters was presented in Holzmüller et al. 2024. Inspired by the remarkable performance of decision trees ensemble methods on tabular data (Shavitt et al. 2018, Grinsztajn et al. 2022, McElfresh et al. 2024), studies have attempted to implement neural network-based decision forest by ensuring differentiability (Popov et at. 2019, Marton et al. 2024). More recently, neural networks built upon the self-attention mechanism of Transformer have been proposed (Padhi et al. 2021, Gorishniy et al. 2021, Hollmann et al. 2022, Huang et al. 2020 and Arik et al. 2021).

However, given that deep learning has also shown to fare poorly with sub-par performance on structured tabular data (Borisov et al. 2022, Shwartz-Ziv et al. 2022), it is not well suited for omics data. In this work, we propose domain-specific prompt engineering of a generative LLM to linearize tabular pharmacogenomics data. Consequently, we were able to outperform the performances of several state-of-the-art deep learning baselines on the DSP task.

### Supplementary Note 2: Descriptions of Pharmacogenomics Datasets

#### GDSC

The Genomics of Drugs Sensitivity in Cancer (GDSC) (Yang et al. 2012) is an open-source database consisting of screening response data of tumoral cell lines to anticancer treatments. In phase 1 (GDSC1), the sensitivity of 987 cancer cell lines to 320 compounds were assayed, while phase 2 experiment (GDSC2) assayed an additional 809 cancer cell lines to 175 compounds (with some overlapping samples in the GDSC1 assay). Cancer cell lines are characterized by genetic features, such as the mutation state. We downloaded the raw dose-response data for GDSC2 from http://www.cancerrxgene.org/downloads/.

#### CCLE

The Cancer cell line Encyclopedia (CCLE) (Barretina et al. 2012) contains a large-scale genomic data (e.g., gene expression) obtained using Affymetrix U133 + 2 arrays for 947 human cancer cell lines and response data for around 500 of the cell lines to 24 drug compounds across 36 tumor types. There are 491 common cancer cell lines having both drug sensitivity measurements and gene expression profile data. The data were downloaded from the CCLE website (http://www.broadinstitute.org/ccle) and PharmacoGx R package.

#### DrugComb

The DrugComb (Zagidullin et al. 2019, Zheng et al. 2021) dataset includes data on synergy and sensitivity of drug combinations. It also includes single drug sensitivity that is characterized as a dose-response curve in terms of IC50 with 717,684 single drug screenings from 37 studies (March 2021). For each drug-drug sample in the dataset, we pair each drug with the cell line to form two separate drug-cell line pairs and include the corresponding single drug IC50 scores. The data was downloaded from https://drugcomb.org/.

#### PRISM

The secondary PRISM Repurposing dataset (Corsello et al. 2020) is a drug repurposing database that includes results of pooled-cell line chemical-perturbation viability screens for 1448 drug compounds screened against 499 cell lines. The original clinical indications for majority (53%) of the active compounds were for non-oncology purposes. The data was downloaded from https://depmap.org/portal.

### Supplementary Note 3: Cancer Cohorts

We focus on the following five cancer tissue types in our evaluation - *lung*, *thyroid*, *breast*, *brain*, *colon/stomach* - as they are available across all four datasets (GDSC, CCLE, DrugComb, PRISM) for evaluation. Note that the aforementioned tissue names could be different in some datasets (e.g., brain, central nervous system). We removed the drug-cell line pairs with missing drug or cell line from each cohort dataset which resulted in the dataset sizes shown in **Supplementary Table 1**. For training and testing GPT, we randomly divided each dataset with an 80%-20% stratified split. Note that for GDSC, as the class distributions are imbalanced with fewer drug-cell line pairs available for the sensitive class (<15%), as shown in **Supplementary Table 2**, we oversampled the minority class (i.e., sensitive) in the training sets of the tissues in GDSC.

### Supplementary Note 4: Comparative Evaluation between GPT-3, GPT-3.5, GPT-4

We carried out fine-tuning analyses on CCLE’s tissue cohorts using the GPT-3, GPT-3.5 and GPT-4 models. This entailed sending a fine-tuning job request to OpenAI’s Completions API for GPT-3. For GPT-3.5 and GPT-4, we sent the respective fine-tuning job requests to OpenAI’s Chat Completions API, with each drug-cell line input in the dataset formatted as a conversation (**Supplementary Figure 1**). We found that GPT-3.5 and GPT-4 performed the same as GPT-3 with no additional improvements; hence this positions GPT-3 as a more cost-effective option for drug sensitivity prediction.

### Supplementary Note 5: Learning Approach

We employ four different paradigms to adapt GPT-3 for downstream drug sensitivity prediction. (i) *Fine-tuning* is based on supervised learning where the pretrained GPT-3 model is trained on prompt-completion pairs in the training set for 4 epochs and its performance is evaluated on the test set. (ii) *Zero-shot* relies on the unsupervised pretrained GPT-3 model’s parametric knowledge to generate drug sensitivity response during inference. This is facilitated by inputting textual prompt that includes the test sample and task-specific description. (iii) *Few-shot* is similar to zero-shot but also inserts a handful of demonstration examples (“shots”) into the prompt as domain-specific knowledge to learn from, referred to as in-context learning. For zero-shot and few-shot experiments, we applied GPT-3 only on the test split for evaluation. (iv) We introduce a weakly supervised learning approach, *clustering embeddings* (Fei et al. 2022), that utilizes text embeddings. These embeddings are first retrieved from OpenAI’s embedding API by feeding drug-cell line prompts as the input. The retrieved embeddings are then clustered using a Bayesian Gaussian Mixture Model (BGMM) (Bishop C 2006). The clusters are initialized using pseudo-labels derived from the fine-tuning predictions and iteratively refined using BGMM. We compare the final cluster assignments for the test samples against their ground truths for evaluation.

We performed each evaluation 3 times and considered the average as the final F1 score.

### Supplementary Note 6: Feature Descriptions

The first feature is the drug’s molecular structure information (MS) in the simplified molecular input line entry specification (SMILES) format (Weininger D. 1998). We use additional molecular or genomic context (MGC) as the second feature and this feature varies across the datasets due to the lack of another common feature among all four datasets. In particular, it includes *gene mutation* in segment for GDSC, *gene expression* for CCLE, *drug-drug synergy* information for DrugComb and *drug’s mechanism of action* (*MOA*) for PRISM. For CCLE, the gene expression data of a cell line is available for about 20,000 genes, corresponding to a vector of the same length. As a result of the high dimensionality overhead of the gene expression data, we sort the gene vector for each cell line by their transcript level values in descending order and select the top 200 genes to represent the gene expression of the cell line as the MGC feature. For DrugComb, we use a Loewe score greater than 5 to categorize the drug-drug synergy information as either ‘synergistic’ or ‘not synergistic’ (Li et al. 2023).

Examples of input prompt templates with MGC are depicted in **Supplementary Figures 3A-3D** for the four datasets. These two features are integrated with the drug-cell line input pair (Basic Information (BI)), leading to three feature groups in total (BI + MS, BI + MGC, BI + MS + MGC), that are evaluated per dataset.

### Supplementary Note 7: Analysis of the drug-pathway associations

When a drug exerts its effects on a cell line, it is known to affect a related pathway rather than a single target (Wang et al. 2021). Henceforth, to evaluate the interpretability of GPT-3, we identify the associations between drugs and signaling pathways, with the possibility to obtain biological insights. In order to infer the drug-pathway associations, we generate the heatmap of Pearson correlations between the predicted drug responses (IC50) and pathway activity scores. However, as large language models such as GPT-3 are naturally suited for classification rather than regression problems, we could not use GPT-3 to predict the IC50 scores. So as a work around, we leverage the embedding model available through OpenAI’s embedding API (https://api.openai.com/v1/embeddings) to retrieve text embeddings of the input prompts in the CCLE dataset. The input prompts include the drug name, cell line name and top 200 genes in the gene expression from the CCLE dataset. We then train a Random Forest model using the GPT-3-derived embeddings as the input features and optimize it to predict the IC50 score. While to calculate the pathway activity scores, we consider the following 11 cancer-relevant pathways from PROGENy (Schubert et al. 2018): *EGFR*, *MAPK*, *PI3K*, *VEGF*, *JAK-STAT*, *TGFb*, *TNFa*, *NFkB*, *Hypoxia*, *p53-mediated DNA damage response*, and *Trail*. Subsequently, PROGENy is applied to the gene expression data from the CCLE dataset to calculate the pathway activity score of each cell line.

In the computed Pearson correlations between the predicted drug responses (IC50) and pathway activity scores of the cell lines (**Figure 6A(i)**), a negative correlation indicates that the drug inhibits the pathway by reducing the expression of genes in the pathway (i.e., assistant), while a positive correlation denotes pathway activation induced by the drug via increasing the expression of genes in the pathway (i.e., resistant) (Wang et al. 2021). These inferred drug-pathway associations are compared side by side to the actual drug-pathway associations (**Figure 6A(ii)**) - obtained by computing Pearson correlations in relation to the ground truth drug responses in the CCLE dataset. We quantify the extent of similarity between the two distributions by performing the Kolmogorov-Smirnov test between the predicted and actual Pearson correlations. The p-values measured by the test based on the null hypothesis that the two distributions are identical (p-value > 0.05) are annotated in red in **Figure 6A**.

### Supplementary Note 8: Baseline Models

For baseline comparisons, we consider the following Transformer-based pretrained language models: (i) *BERT* (Devlin et al. 2018) is a masked language model that leverages Transformer’s encoder; (ii) *BART* (Lewis et al. 2019) is a generative language model that leverages a BERT-like encoder and GPT-like decoder; (iii) *BioBERT* (Lee et al. 2020) is the domain-specific variant of BERT pretrained on large biomedical corpora. (iv) *DistilBERT* (Sanh et al. 2019) is a smaller version of BERT that leverages knowledge distillation during pretraining; (v) *RoBERTa* (Liu et al. 2019) is similar to BERT but is pretrained using only masked language modeling objective with different hyperparameters and (vi) *ALBERT* (Lan et al. 2019) is a parameter-efficient version of BERT enabled through factorized embedding parameterization and parameter sharing across layers. We used the models provided by HuggingFace (https://huggingface.co/). All models are fine-tuned on the training set for four epochs and evaluated on the test set. We also compare against three existing drug response models: (i) *SWNet* (SWN) (Zuo et al. 2021) adopts a dual convergence model architecture to simultaneously encode the drug and genomic features using graph neural network and convolutional neural network, respectively. It then exploits late integration between the two representations before passing through the prediction subnetwork; (ii) *PaccMann* (PM) (Manica et al. 2019) proposes a multimodal architecture and explores various models for encoding drug representation from SMILES, while using an attention-based gene encoder and (iii) *ConsDeepSignaling* (CDS) (Zhang et al. 2021) integrates signaling pathway information with genomic features through a gene-pathway connection matrix and trains a deep belief network. As these three models were optimized for IC50 prediction in the published works, we tweaked the models’ implementations (downloaded from the respective GitHub repository) for reformulation as a binary classification task to maintain a fair comparative analysis with GPT-3. We evaluated these three models on the GDSC dataset.

### Supplementary Note 9: Baseline Comparisons

On the GDSC dataset, GPT-3 asserted superior F1 performance over ALBERT, BART, BERT, BioBERT, DistilBERT, RoBERTa, CDS, SWN, PM with mean performance gains over all tissues of 5.6% (p-value=5e-05), 5.4% (p-value=5e-05), 5.4% (p-value=9e-05), 5.2% (p-value=0.0001), 5.2% (p-value=2.2e-05) and 5.6% (p-value=2.2e-05), 43% (p-value= 0.0004), 9% (p-value=0.06) and 68% (p-value=0.02), respectively. GPT-3’s performance improvements on the other datasets are as follows: CCLE dataset - ALBERT (22%; p-value=0.02), BART (8.7%; p-value=0.24), BERT (16%; p-value=0.09), BioBERT (12%; p-value=0.01), DistilBERT (5.2%; p-value=0.008), and RoBERTa (18%; p-value=0.009); DrugComb dataset - ALBERT (10%; p-value=0.91), BART (3.6%; p-value=0.73), BERT (0.73%; p-value=0.53), BioBERT (7.6%; p-value=0.85), DistilBERT (1.5%; p-value=0.49), and RoBERTa (8.7%; p-value=0.51); and PRISM dataset - ALBERT (6.2%; p-value=0.37), BART (6%; p-value=0.01), BERT (2.3%; p-value=0.02), BioBERT (5%; p-value=0.007), DistilBERT (6.2%; p-value=0.007), and RoBERTa (6.2%; p-value=0.007).

### Supplementary Note 10: Few-Shot Prompting based on Structure Similarity of Drugs

Drugs with similar chemical structure have shown to exhibit similar sensitivity patterns (Zhang et al. 2015). We carried out a subanalysis wherein we probe if exploiting this phenomenon in GPT-3’s in-context learning would refine its predictive power. We first computed the Tanimoto coefficients between the drugs’ SMILES strings to find the structurally similar drug compounds in our dataset, as visualized in **Figure 6B(i)** for 15 drugs from the DrugComb dataset. We then test GPT-3 for 10-shot inference on each drug by providing five ‘sensitive’ in-context examples in the form of drug’s name, cell line’s name and drug’s SMILE, that are associated with the corresponding structurally similar drugs (high Tanimoto score). While five ‘resistant’ examples are also provided but from the not similar drugs (low Tanimoto score). The results are reported in **Figure 6B(ii)** for 13 of the 15 drugs and compared against performance with randomly selected in-context examples (Random); drugs 5-fluorouracil and vorinostat were excluded from the evaluation as they have low Tanimoto scores with the other drugs. The results suggest that prompting GPT-3 with the context of structural similarity acts as inductive bias that could improve its generalization capability, as majority of the drugs (9 out of 13) had boosted performances by including carefully selected demonstration examples - filtered through Tanimoto coefficient rather than randomly sampled - although the differences are not statistically significant (p-value=0.33).

**Supplementary Table 1:**
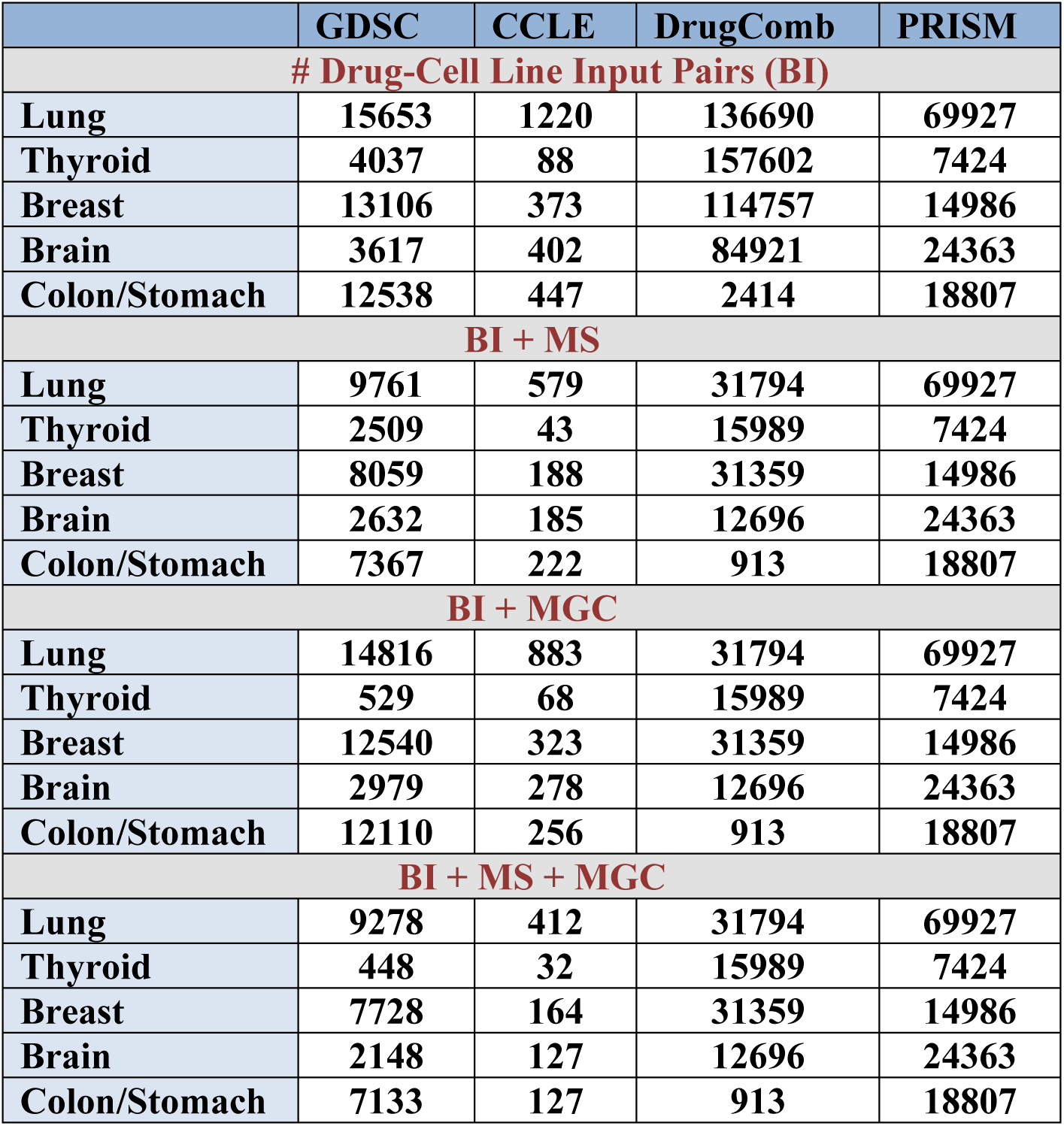
Summary of benchmark pharmacogenomics datasets across tissue types using different feature combinations. We used the following abbreviations: BI (basic information), MS (molecular structure information), MGC (additional molecular or genomic context).

|  | GDSC | CCLE | DrugComb | PRISM |
| --- | --- | --- | --- | --- |
| **# Drug-Cell Line Input Pairs (BI)** |  |  |  |  |
| Lung | 15653 | 1220 | 136690 | 69927 |
| Thyroid | 4037 | 88 | 157602 | 7424 |
| Breast | 13106 | 373 | 114757 | 14986 |
| Brain | 3617 | 402 | 84921 | 24363 |
| Colon/Stomach | 12538 | 447 | 2414 | 18807 |
| **BI + MS** |  |  |  |  |
| Lung | 9761 | 579 | 31794 | 69927 |
| Thyroid | 2509 | 43 | 15989 | 7424 |
| Breast | 8059 | 188 | 31359 | 14986 |
| Brain | 2632 | 185 | 12696 | 24363 |
| Colon/Stomach | 7367 | 222 | 913 | 18807 |
| **BI + MGC** |  |  |  |  |
| Lung | 14816 | 883 | 31794 | 69927 |
| Thyroid | 529 | 68 | 15989 | 7424 |
| Breast | 12540 | 323 | 31359 | 14986 |
| Brain | 2979 | 278 | 12696 | 24363 |
| Colon/Stomach | 12110 | 256 | 913 | 18807 |
| **BI + MS + MGC** |  |  |  |  |
| Lung | 9278 | 412 | 31794 | 69927 |
| Thyroid | 448 | 32 | 15989 | 7424 |
| Breast | 7728 | 164 | 31359 | 14986 |
| Brain | 2148 | 127 | 12696 | 24363 |
| Colon/Stomach | 7133 | 127 | 913 | 18807 |

**Supplementary Table 2:** Class distributions in the training sets of datasets across tissues.

|  |  |  |  |
| --- | --- | --- | --- |
| **GDSC** |  | *Sensitive* | *Resistant* |
|  | **Lung** | 1414 | 11108 |
|  | **Thyroid** | 372 | 2857 |
|  | **Breast** | 1236 | 9248 |
|  | **Brain** | 357 | 2536 |
|  | **Colon/Stomach** | 1237 | 8793 |
| **CCLE** |  | *Sensitive* | *Resistant* |
|  | **Lung** | 177 | 799 |
|  | **Thyroid** | 16 | 54 |
|  | **Breast** | 53 | 245 |
|  | **Brain** | 59 | 262 |
|  | **Colon/Stomach** | 65 | 292 |
| **DrugComb** |  | *Sensitive* | *Resistant* |
|  | **Lung** | 36502 | 72850 |
|  | **Thyroid** | 87928 | 38153 |
|  | **Breast** | 31243 | 60562 |
|  | **Brain** | 24152 | 43784 |
|  | **Colon/Stomach** | 315 | 1616 |
| **PRISM** |  | *Sensitive* | *Resistant* |
|  | **Lung** | 8438 | 47503 |
|  | **Thyroid** | 988 | 4951 |
|  | **Breast** | 1689 | 10299 |
|  | **Brain** | 2890 | 16600 |
|  | **Colon/Stomach** | 2321 | 12724 |

**Supplementary Table 3:** Top three Afatinib treated sensitive and resistant cell lines from the GDSC dataset.

|  | Cell Line | Tissue |
| --- | --- | --- |
| *Sensitive* | MDAMB175VII | Breast |
|  | UACC893 | Breast |
|  | NCIH508 | Stomach |
| *Resistant* | HTCC3 | Thyroid |
|  | NCIH1651 | Lung |
|  | SNUC1 | Stomach |

**Supplementary Figure 1:**
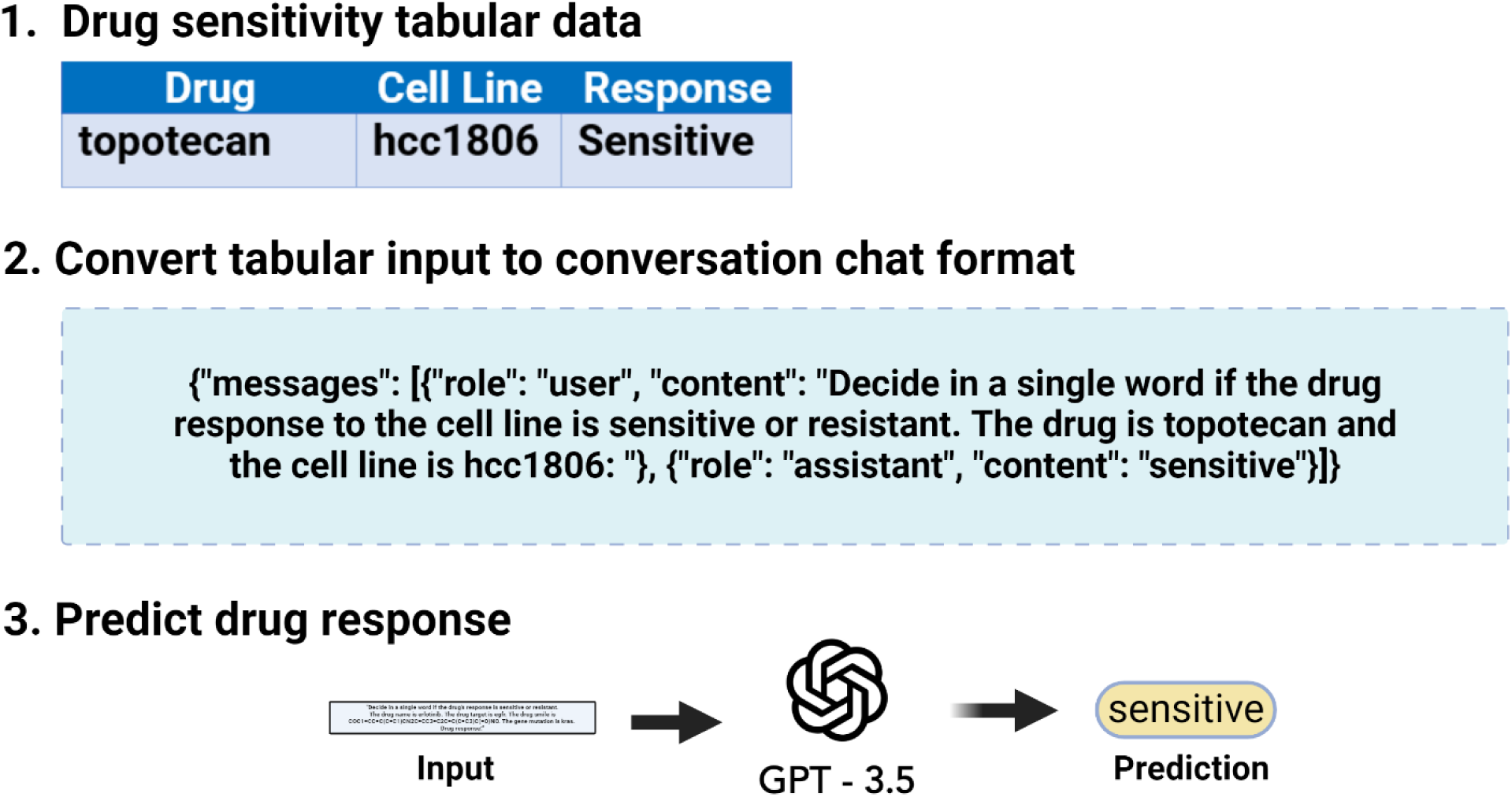
Example demonstrating the conversation chat format for preparing the training samples for fine-tuning with GPT-3.5 and GPT-4.

**Supplementary Figure 2:**
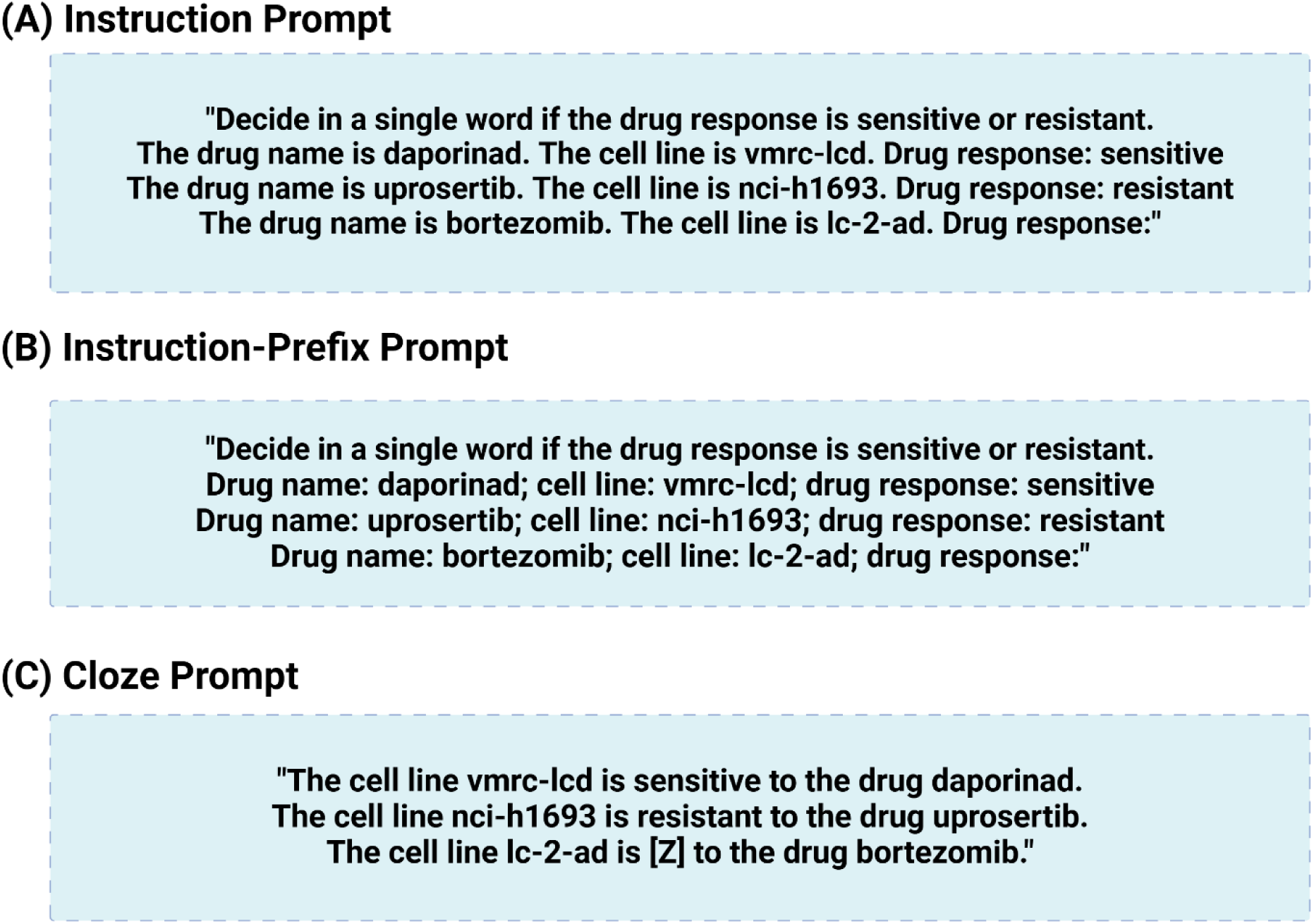
Examples illustrating different prompt templates as input for few-shot evaluation.

**Supplementary Figure 3A:**
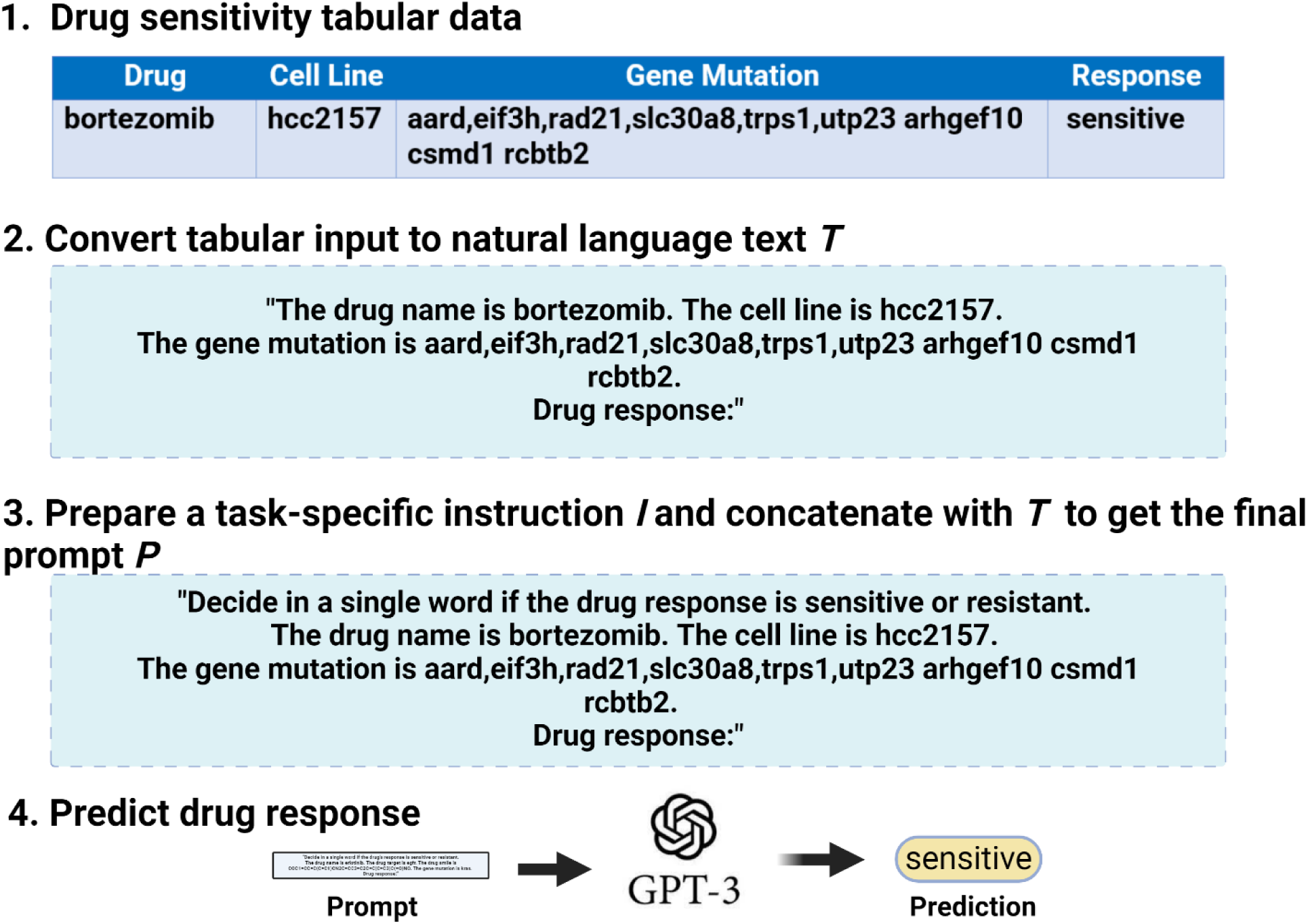
Input prompt example containing feature MGC (BI + MGC) from GDSC dataset.

**Supplementary Figure 3B:**
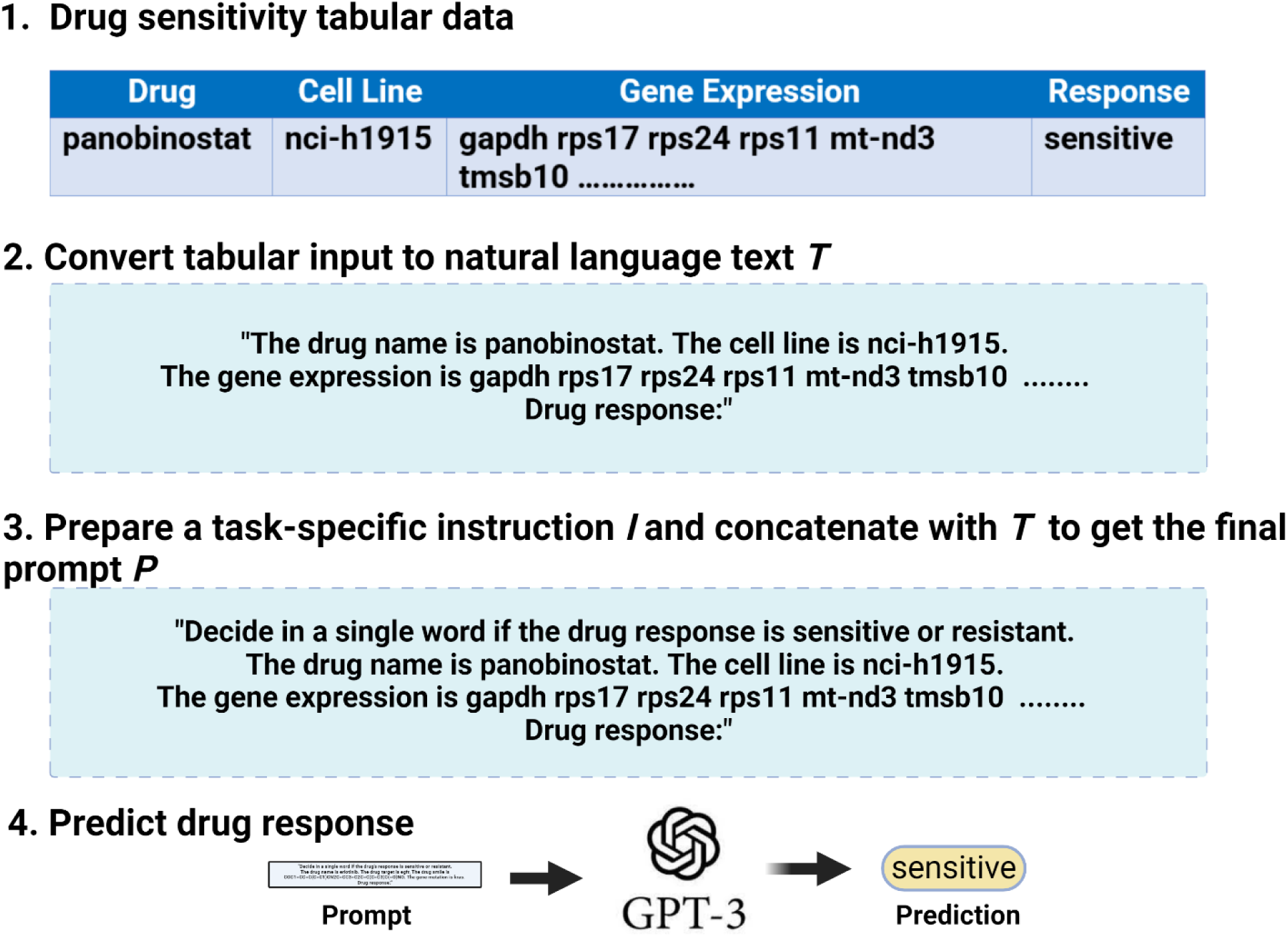
Input prompt example containing feature MGC (BI + MGC) from CCLE dataset. Part of the gene expression is replaced here with ellipsis for brevity.

**Supplementary Figure 3C:**
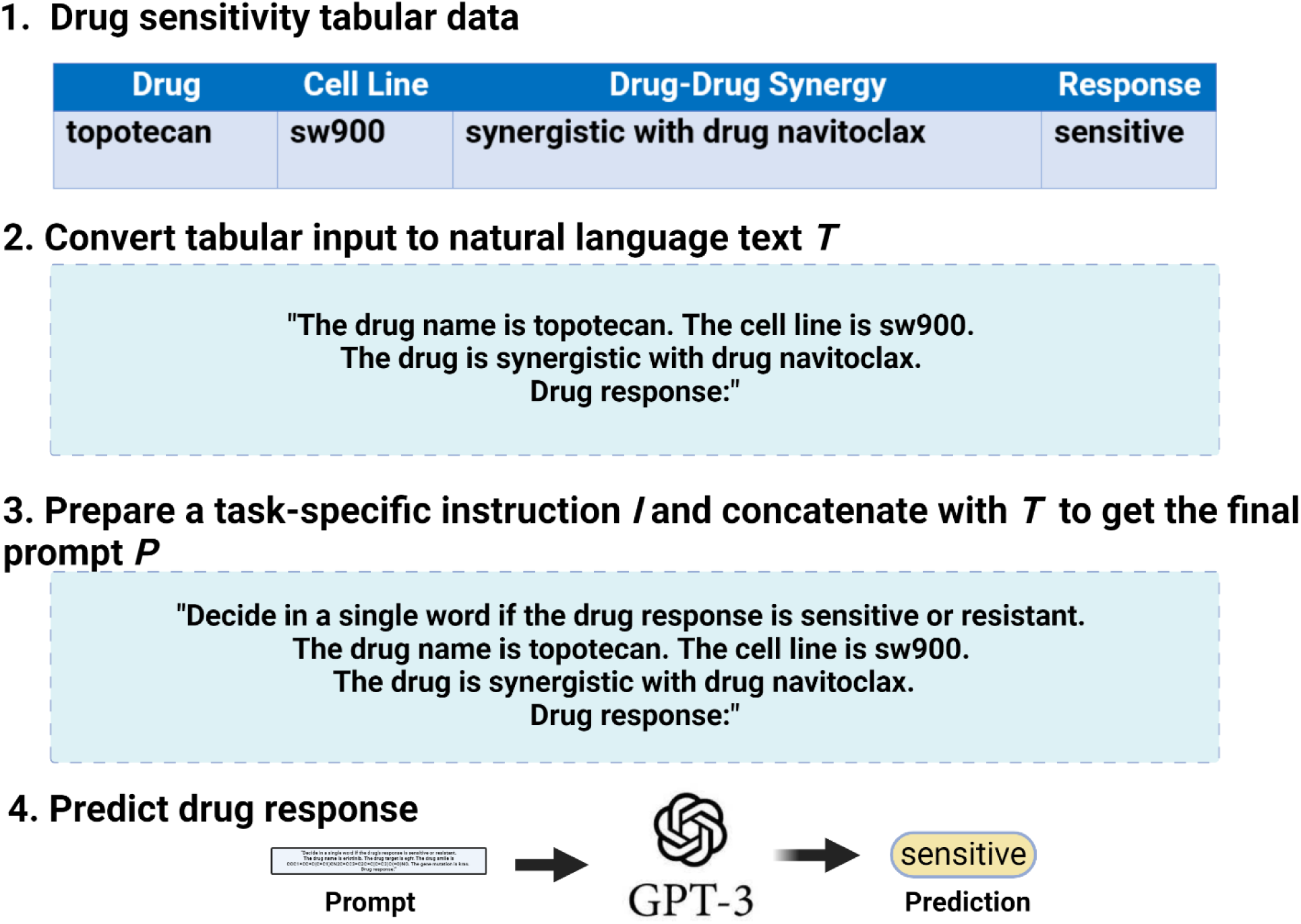
Input prompt example containing feature MGC (BI + MGC) from DrugComb dataset.

**Supplementary Figure 3D:**
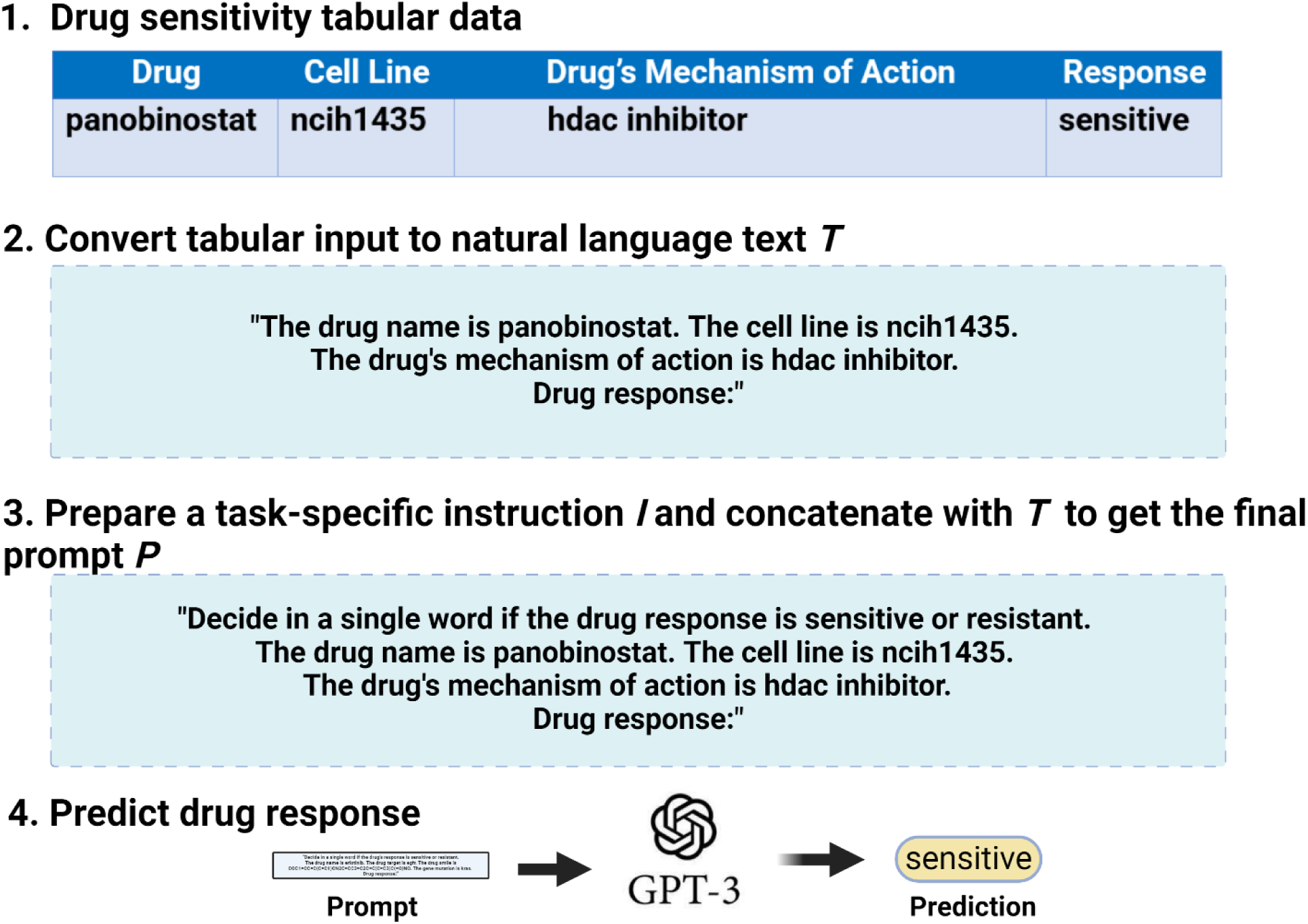
Input prompt example containing feature MGC (Drug + BI + MGC) from PRISM dataset.

**Supplementary Figure 4:**
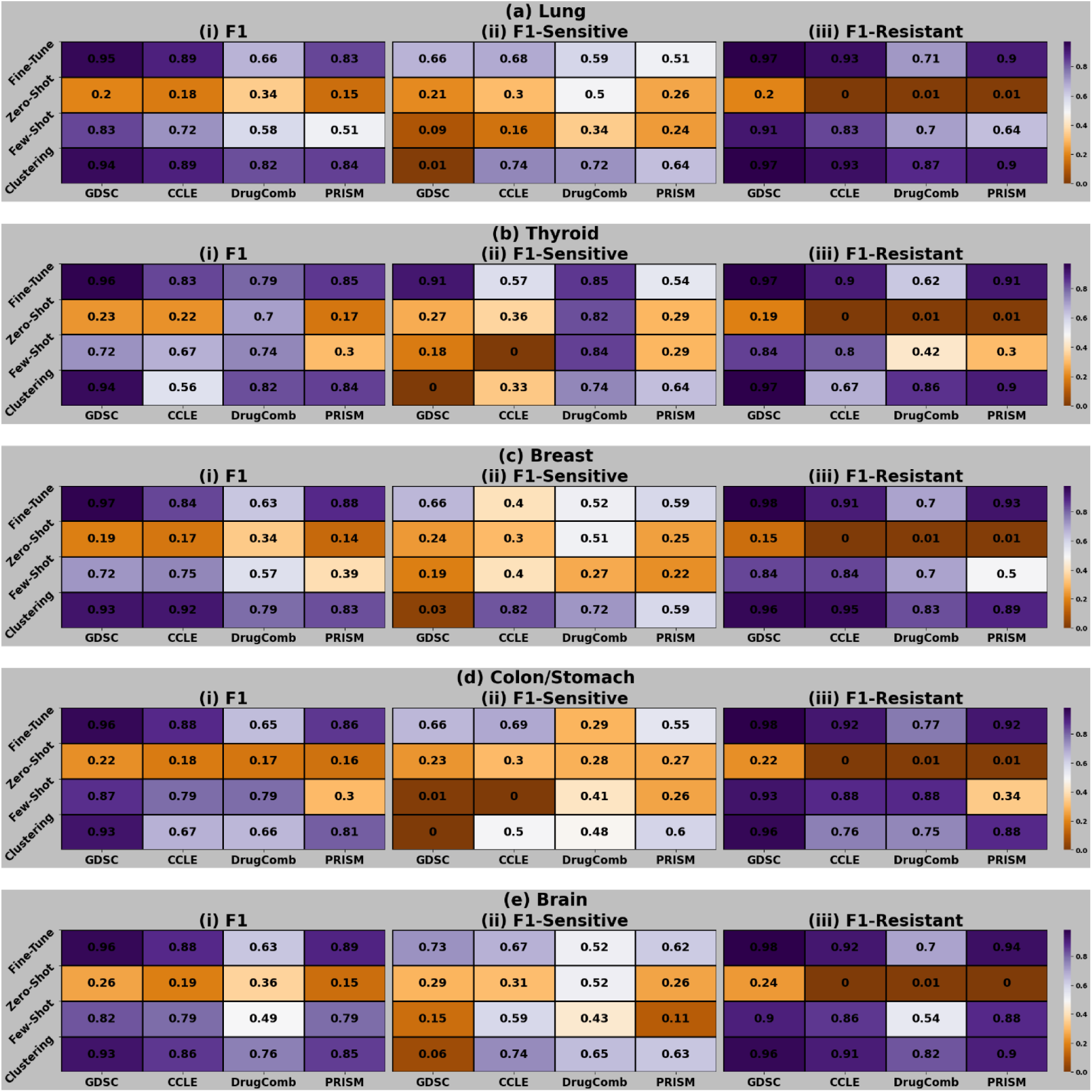
Performance comparisons among learning paradigms across datasets per tissue type.

**Supplementary Figure 5:**
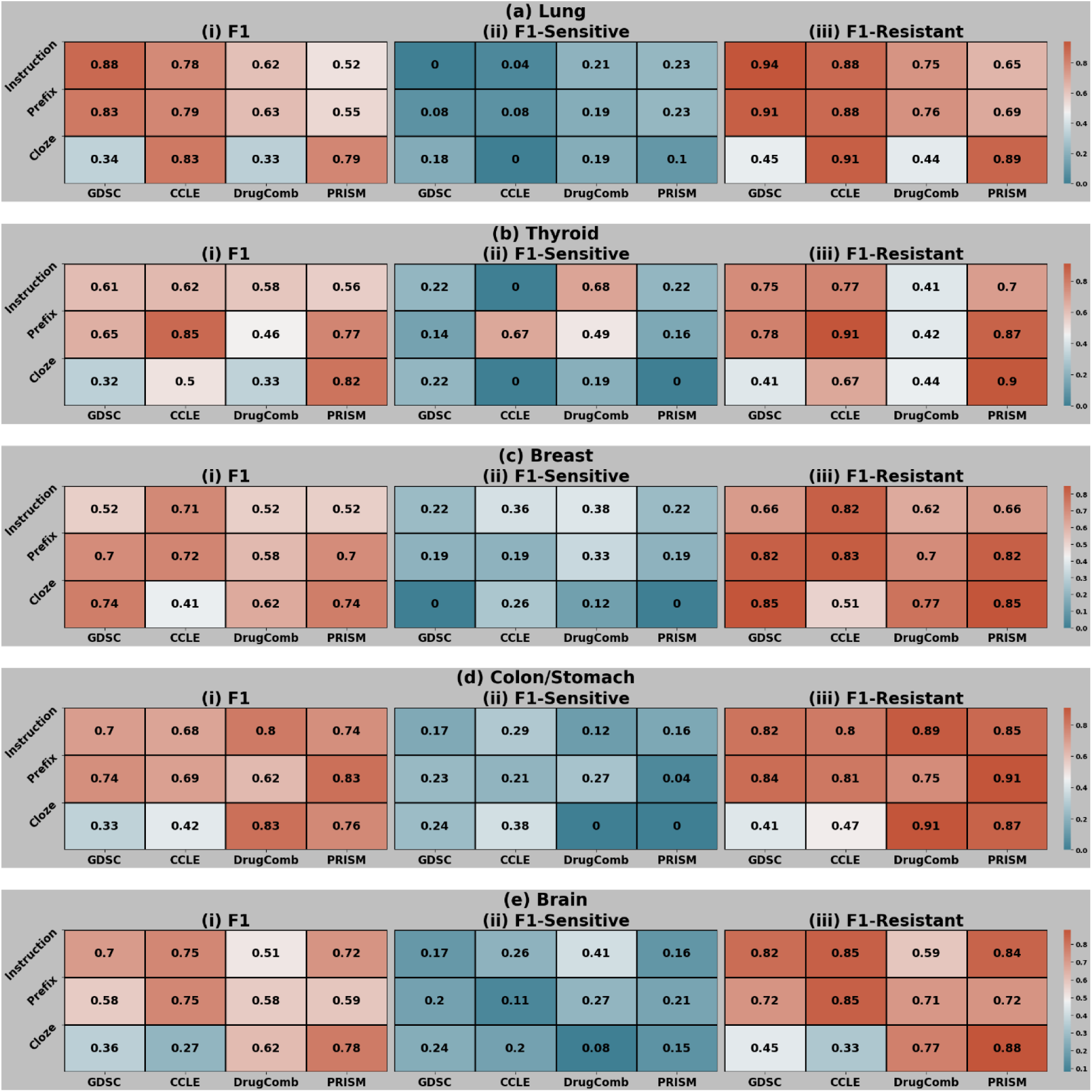
Performance comparisons of prompt templates across datasets per tissue type.

**Supplementary Figure 6:**
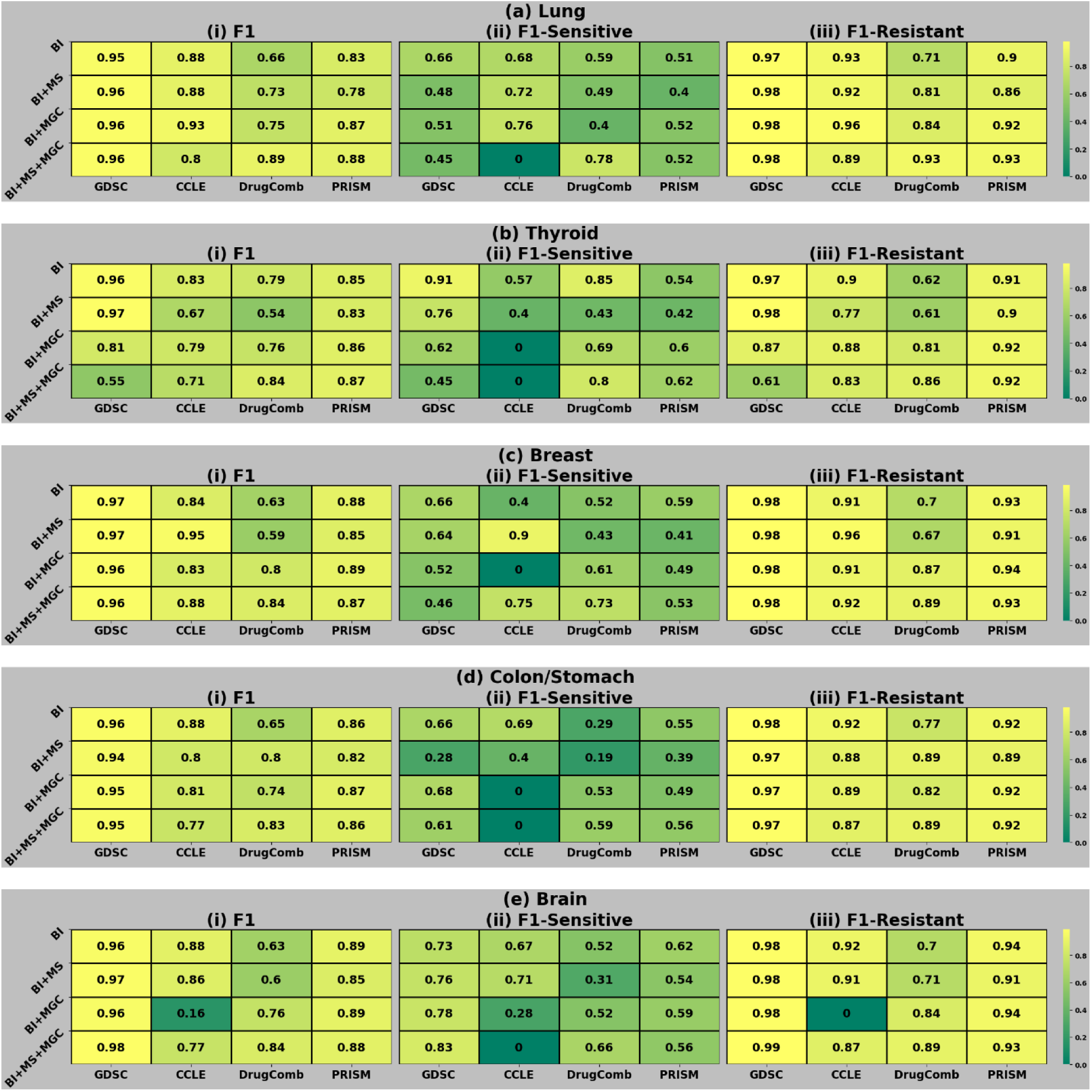
Performance evaluation on feature combinations per tissue type across datasets.

**Supplementary Figure 7:**
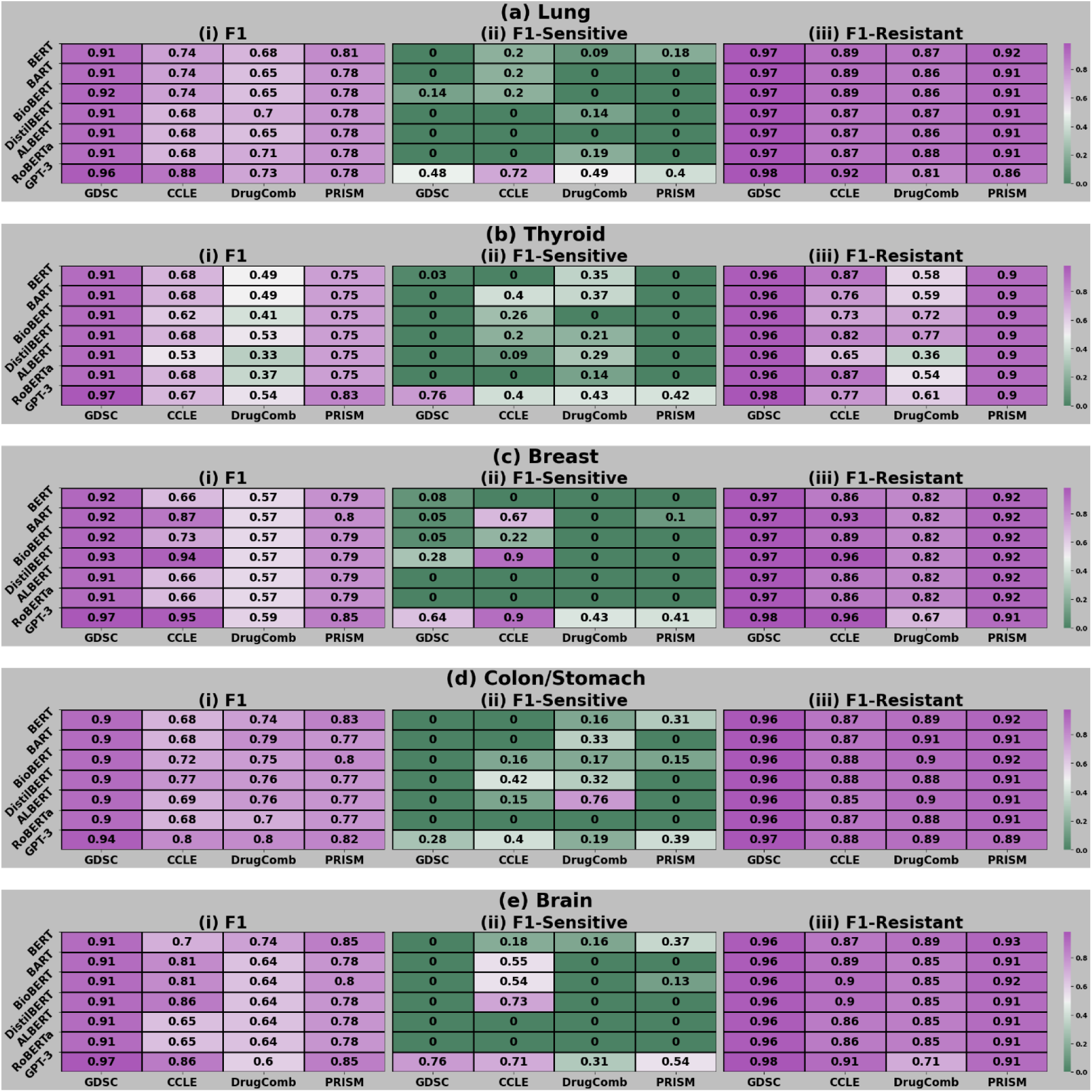
Baseline performance comparisons of GPT-3 with other language models.

**Supplementary Figure 8:**
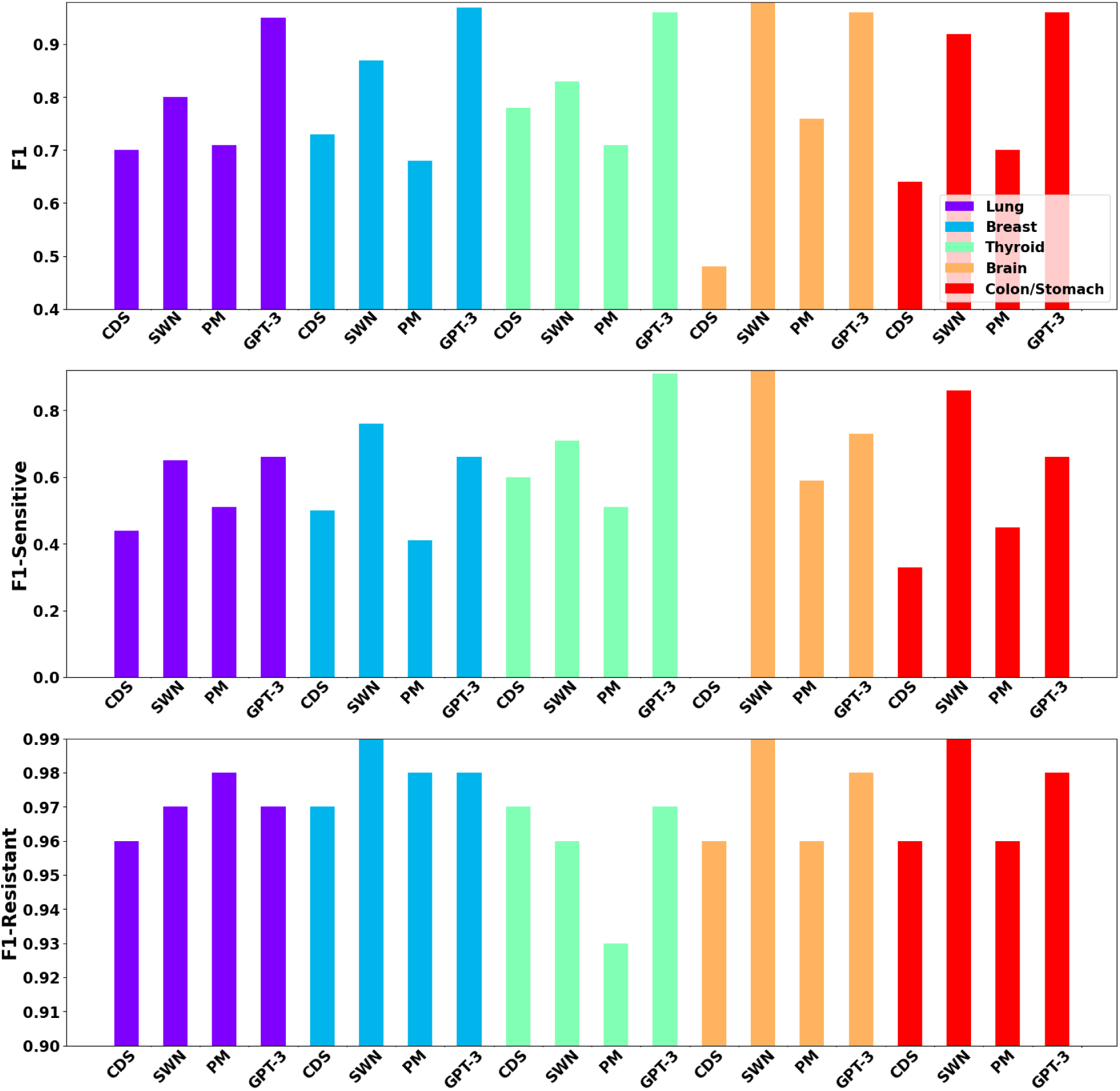
Baseline performance comparisons of GPT-3 with other drug response models.

